# Specialized Molecular Pathways Drive the Formation of Light-Scattering Assemblies in Leucophores

**DOI:** 10.1101/2025.01.12.632588

**Authors:** Y. Barzilay, Z. Eyal, Y. Noy, N. Varsano, T. Olender, S. Bera, T. Lerer-Goldshtein, M. Kedmi, Z. Porat, I. Pinkas, S. Levin-Zaidman, N. Dezorella, D. Gur

## Abstract

Pigmentation plays a vital role in the survival of organisms, supporting functions such as camouflage, communication, and mate attraction. In vertebrates, these functions are mediated by specialized pigment cells known as chromatophores of which, uric acid crystal-forming leucophores remain the least understood, with little known about their molecular mechanisms. A key question in pigment cell biology is whether different crystal chemistries require distinct molecular pathways, or whether similar cellular processes drive the formation of diverse crystals. This study was designed to unravel the uncharacterized process of uric acid crystallization in leucophores and compare them to guanine crystal formation in iridophores and pterin formation in xanthophores. The results of our transcriptomic, ultrastructural, and metabolomic analyses, demonstrate that leucophores share molecular pathways with iridophores, particularly those connected to organelle organization and purine metabolism, but express discrete genes involved in uric acid biosynthesis and storage. Additionally, leucophores share intracellular trafficking and pterin biosynthesis genes with xanthophores, suggesting universally conserved processes. Ultrastructural studies reveal star-like fibrous structures in leucosomes, which likely serve as scaffolds for unique one-dimensional uric acid assemblies that radiate from the core and act as efficient light scatterers. These findings provide new insights into leucophore cell biology and the specialized mechanisms driving molecular crystalline assembly, and reveal that while some cellular processes are conserved, the specific chemistry of each crystal type drives the evolution of distinct molecular pathways.

## Introduction

Pigmentation in organisms is essential for survival, enabling camouflage, vision, social signaling, and mate attraction^1-5^. In vertebrates, these diverse functions are mediated by specialized pigment cells derived from neural crest cells (NCCs)^6-8^. Mammalian and avian pigmentation is controlled by a single cell type, the melanocyte^9-11^. However, poikilothermic vertebrates, such as teleost fish, possess a broader range of pigment cells known as chromatophores^12-15^. These chromatophores can be classified into light absorbers, like black or brown melanophores (M), yellow to red xanthophores (X), and cells that reflect or scatter light, like iridescent iridophores (I) and white leucophores (L)^12,13,16,17^. Pigment formation in all chromatophores occurs within specialized membrane-bound organelles—melanosomes in M, pterinosomes in X, iridosomes in I, and leucosomes in L—all of which are thought to belong to the lysosome-related organelle (LRO) family^18-22^. Among teleost species, medaka (*Oryzias latipes*) has all four types of pigment cells, including leucophores^23,24^, which remain the least understood in terms of their molecular mechanisms.

The ability of different chromatophores to manipulate light by reflection, scattering, or absorption, not only generates the visual effects critical for survival but also supports physiological and ecological functions. The observed diversity of chromatophores enables the formation of complex and dynamic coloration patterns, and made them powerful models for studying cell fate specification, differentiation, and the evolution of pigmentation.

While guanine crystals are widely recognized as the most abundant molecular crystals used by organisms^1,25-29^, uric acid assemblies are prevalent across various functional biological systems and are commonly associated with excretion in many species^30-35^. In humans, the uncontrolled formation of uric acid crystals can lead to conditions such as kidney stones and gout, which affect millions worldwide^36-39^. In contrast, organisms like medaka tightly regulate uric acid crystal formation and dissolution within leucophores.

Leucophores are particularly intriguing chromatophores due to their dual characteristics. Traditionally, these cells were grouped with iridophores, because both cell types produce membrane bound purine-based crystals that optically interact with light: uric acid in leucophores and guanine in iridophores^40,41^. However, recent studies suggest that leucophores may also share developmental and genetic similarities with xanthophores^42-44^. This duality raises important questions about the origins, molecular mechanisms, and evolution of leucophores, particularly in species like medaka, where they represent a prominent feature of the fish phenotype^14^.

Recent work has elucidated the molecular mechanisms responsible for guanine crystal formation in iridophores^19,45-47^. Therefore, investigating the processes that drive uric acid crystal formation in leucophores now offers a unique opportunity to compare these two systems. This comparison raises key questions as to whether leucophores and iridophores employ similar cellular processes for crystal formation, or whether the distinct chemistries of guanine and uric acid require different molecular pathways. Furthermore, extending this comparison to xanthophores, which produce pteridine and carotenoid-based pigments, adds an additional layer of information about the relationships between the chemistry and the cellular processes that facilitate and regulate pigment formation and function.

This study was designed to identify the cellular and molecular mechanisms governing uric acid crystal formation in leucophores, while explore their similarities and differences with guanine crystal formation in iridophores and pterin and carotenoid production in xanthophores, using medaka as a model system. A combination of transcriptomic and ultrastructural analyses, including single-cell RNA sequencing (scRNA-seq), advanced electron microscopy, and spectroscopy were employed to identify the biosynthesis, micro-environment, accumulation, and crystallization pathways in leucophores, and compare them with those in iridophores and xanthophores. The aim was to determine whether similar intracellular molecular mechanisms can drive the formation of chemically distinct assemblies across pigment cell types.

Our findings reveal that while several molecular pathways involved in uric acid formation in leucophores overlap with those driving guanine formation in iridophores, leucophores also express unique gene sets specific to uric acid biosynthesis, storage, and accumulation. Additionally, leucophores and xanthophores share a subset of genes related to intracellular trafficking and pterin biosynthesis, thereby providing evidence for conserved processes common to these two cell types. Notably, high-resolution cryo-electron microscopy (cryoEM) and micro-Raman spectroscopy revealed that the leucosome structures consist of one-dimensional uric acid assemblies and act as efficient light scatterers, thereby challenging previous assumptions that leucophores contained fully crystalline uric acid.

The results of this study advance our understanding of leucophore function at both the cellular and molecular levels and provide new insights into the common and specialized mechanisms driving the formation of molecular crystalline assemblies with distinct chemistries. The dual nature of leucophores, which share certain characteristics with iridophores and others with xanthophores, illustrates the complexity of pigment cell differentiation and highlights the evolutionary processes that contribute to the remarkable diversity of coloration in vertebrates. In this context a smaller set of upregulated genes was found to be common to all three pigment cells, suggesting that some mechanisms are universally conserved across leucophores, xanthophores, and iridophores.

## Results

### Optical Properties and Cell Isolation

In the absence of transgenic fluorescent lines in medaka, that can be used to specifically label and sort leucophores with a fluorescence activated cell sorter (FACS), we resorted to isolating the cells based on their inherent optical properties. Analysis of the autofluorescence and light-reflecting properties of the different pigment cells using live confocal light microscopy (**Fig. 1**) revealed that while both crystal-containing leucophores and iridophores are visible under incident light and cross-polarized microscopy, only leucophores exhibit strong autofluorescence under multiple laser wavelengths (**Fig. 1C**). In contrast, xanthophores were only visible using the green laser and were not seen by incident light and cross-polarized microscopy. These distinct optical properties are driven by the scattering in leucophores, the reflective surfaces in iridophores, and the light absorption in xanthophores, and highlight the unique adaptations of each chromatophore type. We were able to exploit these distinct optical properties to isolate three different populations of pigment cells: iridophores, leucophores, and xanthophores (**Fig. S1**).

**Figure 1.**
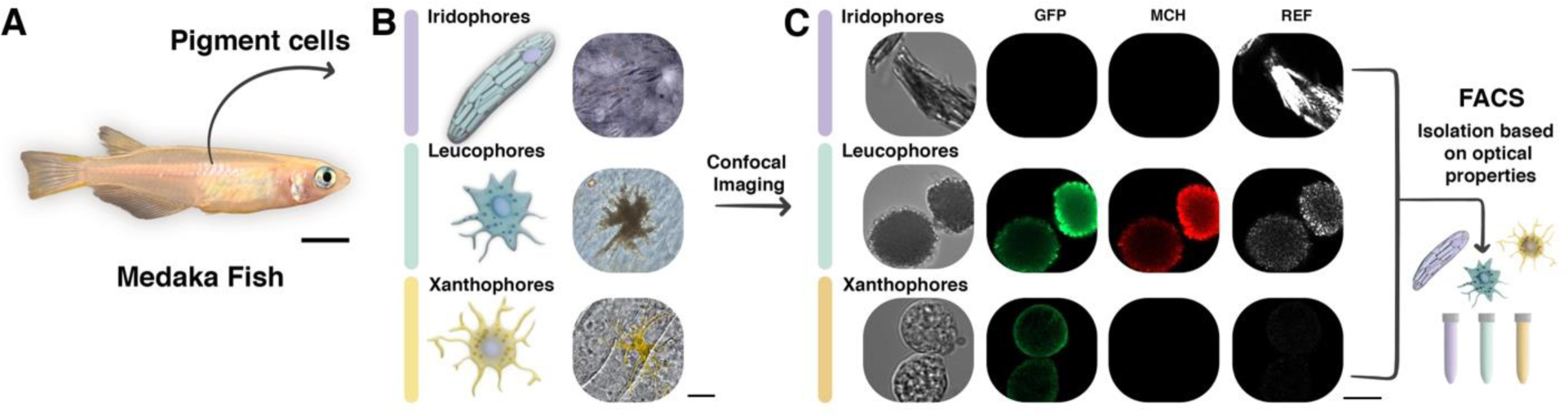
Isolation of different pigment cells in medaka based on their optical properties. **(A)** An adult medaka fish. **(B)** Illustration (left) and bright-field optical microscopy image (right) showing the different pigment cells in medaka. **(C)** Confocal live microscopy of various pigment cells highlighting their distinct optical properties, which were used to isolate them by FACS. Scale bars: A – 5 mm; B – 10 μm; C – 10 μm.

### Ultrastructure of Pigment Cells

As the next step, we compared the ultrastructure of leucophores, xanthophores, and iridophores using electron microscopy (**Fig. 2**). Traditional transmission electron microscopy (TEM) methodologies, dissolved both the uric acid crystals within leucosomes and the guanine crystals within iridosomes, but only minimally effected the content of the pterinosomes. (**Fig. 2 B,G,L**). However, unlike the iridosomes which then appear as white holes, dissolving the uric acid crystals revealed highly stained intricate, star-shaped structures that are probably composed of proteins or lipids or other macromolecules. A similar fibrous structure was recently observed by cryoEM in iridosomes, where parallel fibers were shown to act as a template for nucleation ^46,48^. We then used cryogenic imaging, which allows us to maintain the integrity of the internal crystals and observe the leucophores in as close to their native state as possible. Initially, we examined the different pigment cells *in situ* using cryogenic scanning electron microscopy (cryo-SEM), which enabled us to observe intact round leucosomes (**Fig. 2F**), platelike crystals within iridophores (**Fig. 2A**), and onion-like structures within the pterinosomes, which are the characteristic pigmentary organelles of xanthophores (**Fig. 2K**). The finer details of these organelles and their contents, were imaged in higher resolution by plunge freezing each isolated cell population on a TEM grid and imaging them with cryo-TEM (**Fig. 2 C,H,M**). This method provided a closer look at the organelle content and revealed numerous membrane-bound, neatly arranged concentring rings within the xanthophores (**Fig. 2M**), elongated membrane-bound crystals in the iridosomes (**Fig. 2C**) and, round or star-shaped crystals in the leucosomes (**Fig. 2H**). The 3D morphology of the of the three different organelles was examined further by cryo-electron tomography (cryoET) (**Fig. 2 D,E,I, J, N, O, and Movie S1**). This approach revealed multiple irregular crystals originating from the core of the organelle, possibly draped over the star-like scaffold framework observed by traditional TEM. Examining multiple leucosomes revealed that the morphology of the individual assemblies differs significantly, with each crystal being unique **(Fig. 3)**.

**Figure 2.**
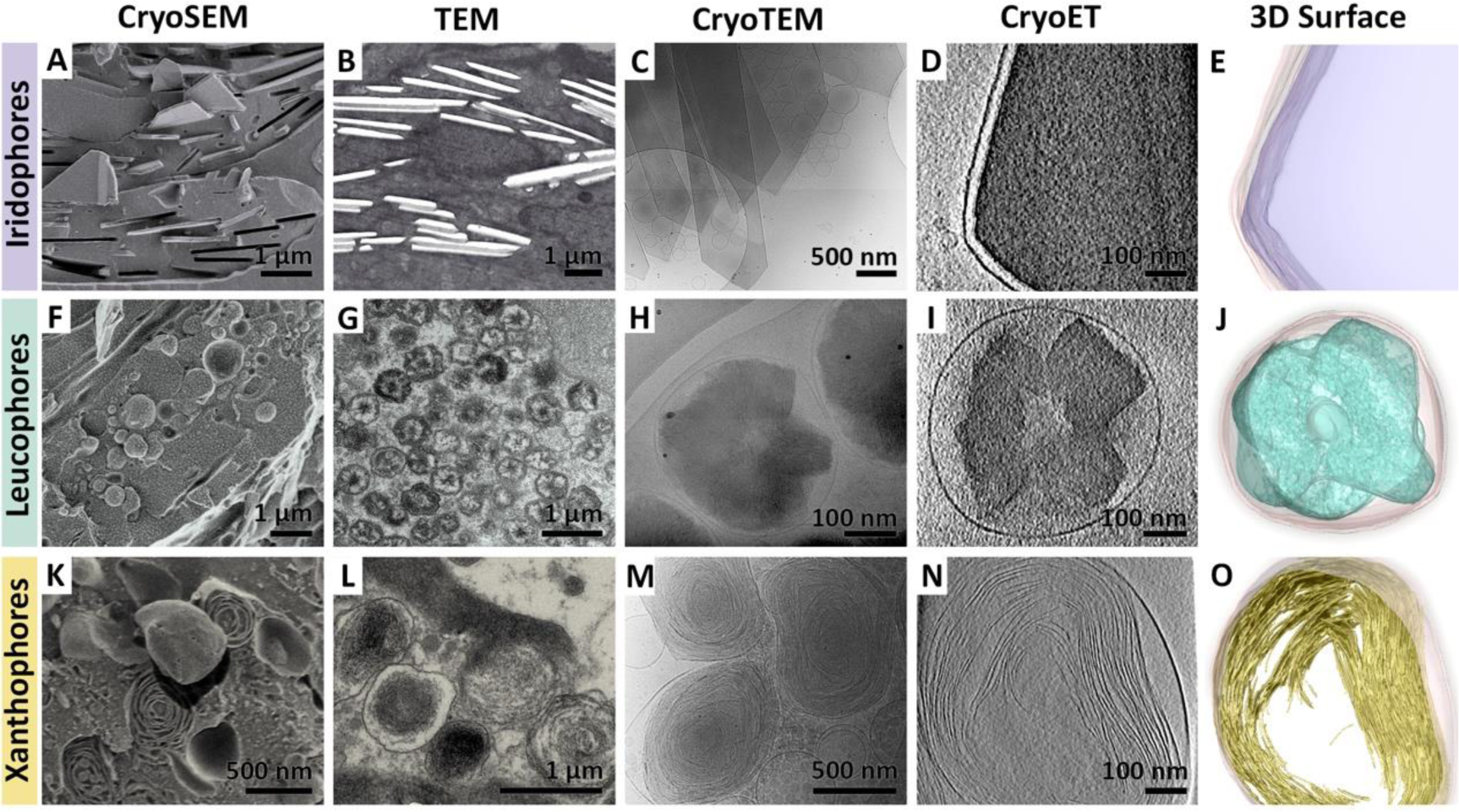
Ultrastructure and organelle morphology of different pigment cells. The ultrastructure of iridophores **(A-E)**, leucophores **(F-J)**, and xanthophores **(K-O)** as observed by various electron microscopy techniques. Iridophores are densely packed with elongated, several microns-in-length, membrane-bound, plate-like guanine crystals. Leucophores are populated with 500-1000 nm leucosomes containing star-shaped crystal assemblies, while xanthophores are characterized by 500-1000 nm onion-like pterinosomes.

**Figure 3.**
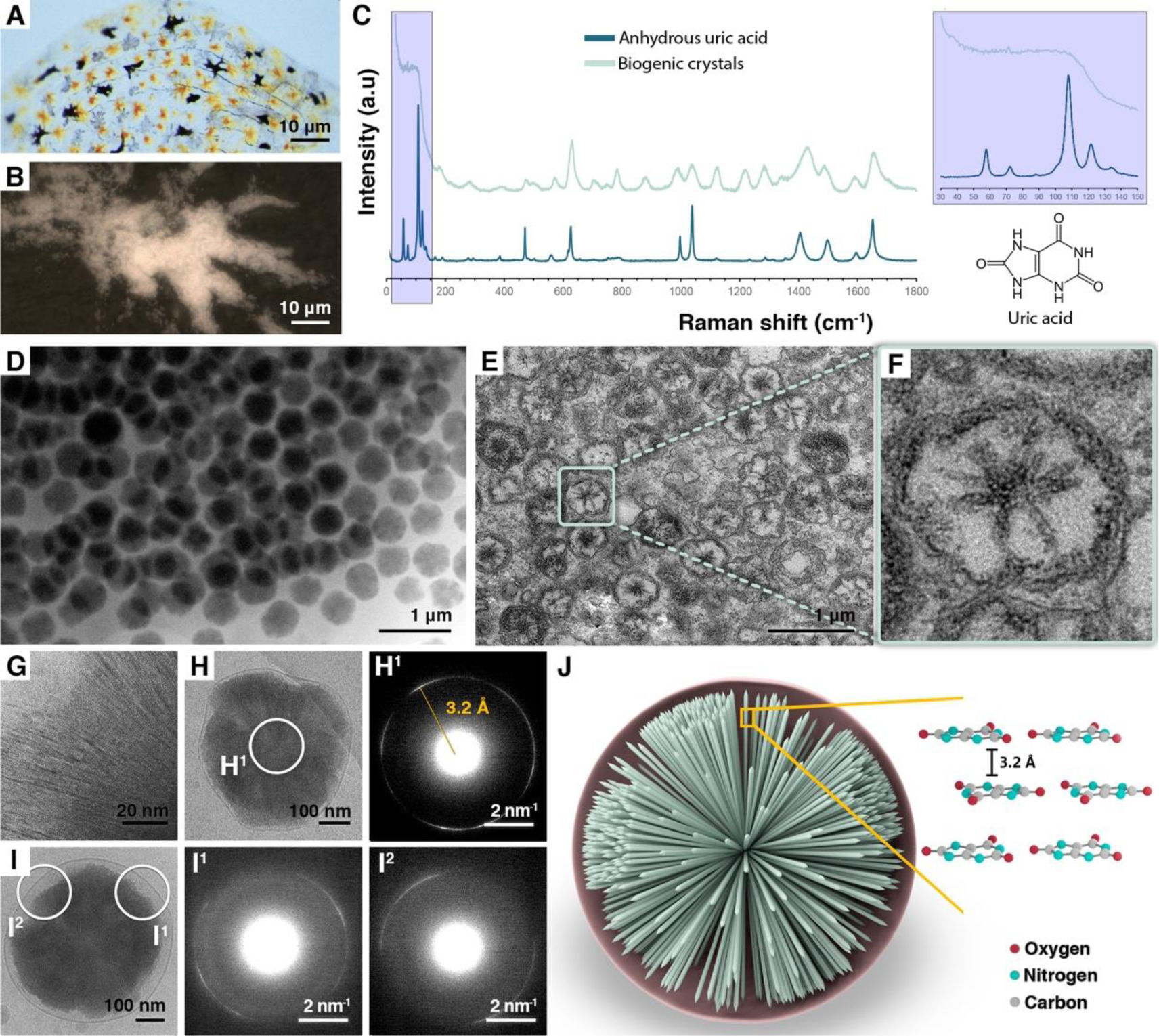
Leucosomes contain a semi-crystalline assembly of uric acid. (**A**) A transmission optical image of a medaka scale, showing different pigment cells. (**B**) A higher magnification view of a leucophore from a medaka scale using incident light (**C**) Micro-Raman spectra of the leucophore showen in panel B, overlaid with the spectra of synthetic uric acid. Inset: Molecular structure of uric acid. Right: low frequency region of the spectrum. (**D**) A cryo-TEM image of a leucophore showing numerous densely packed leucosomes. (**E**) A conventional TEM image of a leucophore, in which the crystalline materials is lost, showing heavily stained fibres in a star-like shape arrangement. (**F**) A higher magnification view of the leucosome marked in (E). (**G**) A high-magnification cryo-TEM image of the interior of an intact leucosome, revealing an arrangement of nanoscopic rods radiating outward from the core of the sphere. **(H-I)** cryo-TEM images and corresponding SAED of an isolated leucosome showing a single diffraction pattern with a spacing of ∼3.2 Å. Collecting an electron diffraction pattern from the center results in a full diffraction ring (H^1^), while moving along the dihedral angles of the particle (I) reveals changes in the reflection angle. This suggests that the stacking axis of the crystal projects radially outward from the center of the leucosome. (**J**) Schematic representation of the sphere showing the spherulitic arrangement of stacked uric acid molecules.

### Chemical Composition of Leucosomes

Leucophores have recently been reported to contain high levels of uric acid, which prompted the, as yet, unconfirmed hypothesis that leucophores contain crystalline uric acid^49^. In order to determine the exact chemical nature of the material within leucophores, we performed *in situ* micro-Raman measurements of leucophores located in medaka scales (**Fig. 3 A-C**). Comparison of the spectra from various reference purine crystals revealed that, despite some differences in spectral features, the particles primarily consist of uric acid (**Fig. 3C and S2**). Notably, unlike a fully crystalline uric acid standard, the Raman spectrum from leucophores lacks the low-frequency peaks typically associated with lattice modes (**Fig. 3C, inset**). This absence suggests that the uric acid molecules within the leucophores do not exhibit the three-dimensional periodicity characteristic of fully crystalline structures. To better understand the structural organization of the uric acid within leucophores, we correlated high-resolution cryoET with electron diffraction (**Fig. 3 G-I**). High-resolution TEM images show that leucosomes contain thin rods, only a few nm in thickness, which are grouped into clusters that project from the core of the leucosome (**Fig. 3G**). The electron-diffraction patterns of whole particles exhibit a single diffraction ring with a spacing of ∼3.20 Å (**Fig. 3H**), which correlates to the pi-stacking direction of the conjugated uric acid molecules^50^. Selected area electron diffraction (SAED) at different locations along the leucosome periphery exhibited diffraction arcs corresponding to the dihedral angle of the leucosome (**Fig. 3I**). Taken together, these data indicate that the uric acid assemblies with the leucosomes are arranged as a spherulite, in which 1D-ordered, stacked assemblies of uric acid project radially outwards from the center of the leucosome (**Fig. 3J**). This arrangement exposes the high refractive index plane of uric acid to light, which optimizes the light-scattering properties of the leucosomes. A similar structure was recently reported for isoxanthopterin crystals in shrimps, where it is responsible for their brilliant white colors^51^.

### Leucophore molecular machinery and pigmentation pathway

Given the distinct differences in ultrastructure and molecular composition, we hypothesized that leucophores would exhibit a unique transcriptomic profile that distinguishes them from xanthophores and iridophores. This was tested by subjecting FACS isolated leucophores, iridophores, and xanthophores to single-cell RNA sequencing (scRNA-seq). Dimensionality reduction and unsupervised clustering surprisingly revealed 13 clusters (**Fig. S3A**). Among these, we identified clusters corresponding to fibroblasts (cluster 3), epithelial cells (cluster 6), and immune cells (cluster 2) that may have been co-isolated inadvertently (**Fig. S4**).

To identify the pigment cells, we searched for known neural crest markers such as *snai1a*, *sox10*, *sox5*, *pax7a* and *ngfrb*^52,53^(**Fig. S5**), which led to the identification of clusters 0, 1, 4, 7, and 12. Of these, the small cluster 12 was designated as iridophores due to the high levels of expression of iridophore markers (e.g., *pnp4a*, *gpnmb*, *alx4b*, *tfec*, *ltk*, *apoda.1*, *fhl3*, **Fig. S6**)^45,54-56^. Similarly, the high levels of expression of genes such as *gch2*, *scarb1*, *plin6*, and *tmem130*, in clusters 0, 1, and 4 lead to their classification as xanthophores (**Fig. S7**)^57-59^. Hereafter, we will refer to these three clusters as the “xanthophore” cluster (**Table S1**).

While cluster 7 also expressed some xanthophore marker genes, including *gch2* and *tmem130*, it also expressed iridophore marker genes such as *pnp4a*, *paics*, and *gmps* (**Fig. S8**). Since leucophores share characteristics with both crystal-forming iridophores and dendritic xanthophores, we suspected that this cluster might represent leucophores. To test this hypothesis, we repeated the FACS and isolated only the population of leucophores, then repeated the scRNA-seq, and reanalysed the entire population, noting whether the cells were isolated in the first round (all three pigment cells) or the second round (only leucophores). While a small percentage of the cells isolated in the second round of FACS were distributed across different clusters, the majority could be assigned to cluster 7, thereby validating this region as the leucophore cluster (**Fig. S3B**).

We compared the expression profiles of the three pigment cell types to uncover shared features and unique distinctions (**Table S2**). The identified leucophore cluster exhibits 797 differentially expressed genes, while the xanthophore and iridophore clusters have 978 and 216 differentially expressed genes, respectively (**Fig. 4B**). To further elucidate the molecular machinery and processes in leucophores, we examined these upregulated genes more closely and compared them to those of iridophores and xanthophores (**Fig. 4 B,C**). The results indicate that leucophores share 44 upregulated genes with iridophores and 285 with xanthophores, whereas iridophores and xanthophores have a significantly lower 25 genes in common (Fisher’s Exact Test: L-X: OR = 15.6, p < 0.001; L-I: OR = 6.8, p < 0.001; I-X: OR = 2.7, p < 0.001). A comparison of the upregulated genes that are common to all three cell types revealed only 18 shared genes (**Table S2**).

**Figure 4.**
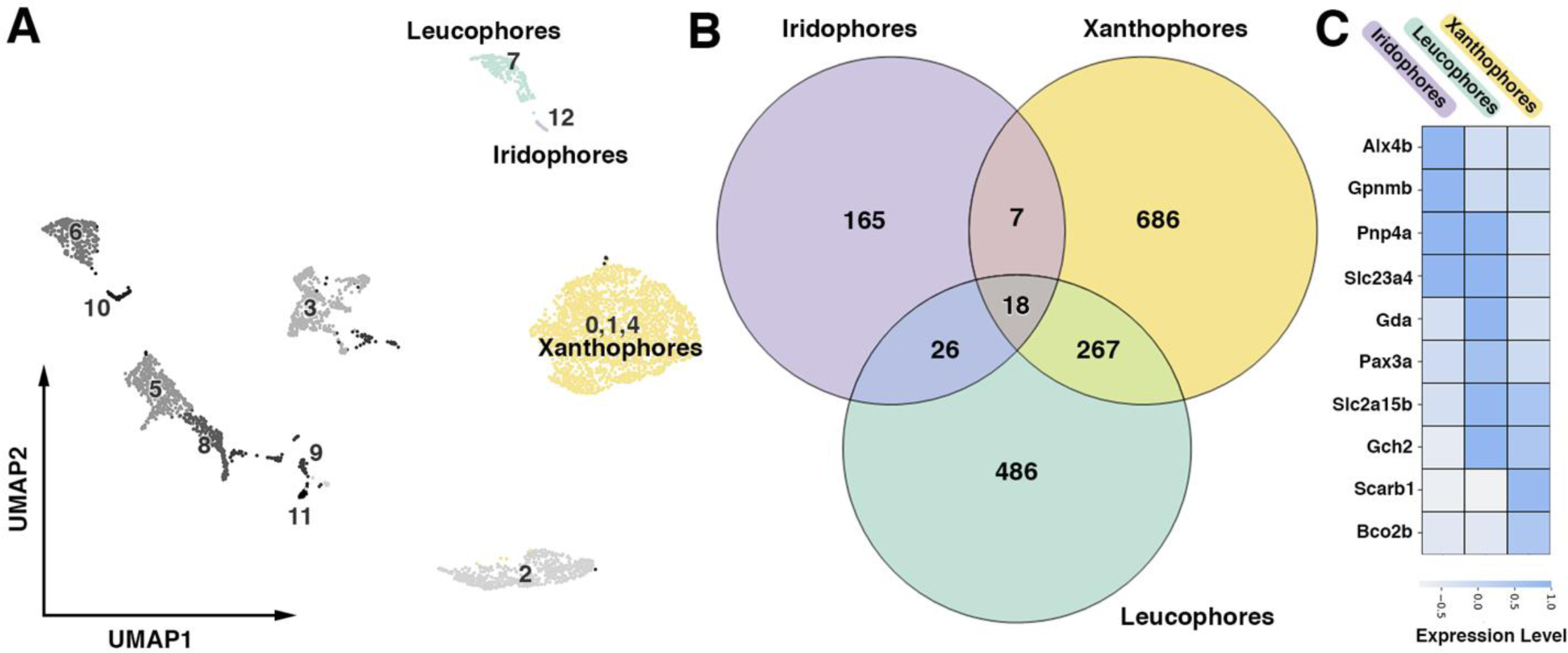
Leucophores share more upregulated genes with both xanthophores and iridophores than xanthophores and iridophores share with each other. **(A)** Two-dimensional UMAP representation of the cells analyzed in the single-cell RNA-sequencing experiment, with pigment cell clusters color-coded. **(B)** Venn diagram illustrating the number of unique and shared genes among iridophores (purple), xanthophores (yellow), and leucophores (turquoise). **(C)** Heatmap showing the average expression levels of selected genes across different pigment cells, showing the overlap between leucophores, iridophores, and xanthophores.

Among the shared genes, iridophores and leucophores both express several solute carrier (SLC) family genes, which are probably involved in nucleobase trafficking and transport^60^, including *slc23a4*, and *slc6a17*. Notably, *slc2a9*, *slc29a4a*, and *slc6a4a* are uniquely upregulated in iridophores, while *slc24a5*, *slc3a2a, slc27a6*, and *slc35G1* are only upregulated in leucophores. Additionally, *slc2a15b* is expressed in both leucophores and xanthophores, whereas *slc2a15a* is common to leucophores xanthophores and iridophores. These findings suggest that while some transport processes are conserved, others are specialized for pigment cell-specific functions, which probably reflects the distinct molecular needs of each cell type.

Similarly, the genes shared by leucophores and xanthophores include *pax7a*, which was recently reported to be involved in the differentiation of both cell types^61^. On the other hand *pax3b* was only upregulated in xantophores, as recently reported^61^. *Csf1ra*, which is known to play a role in xanthophore development^8^ but has not previously been linked to leucophores was also found to be upregulated in leucophores. Other genes common to all three pigment cell types include the lysosomal trafficking regulator gene *lyst^62^*, which has been implicated in the trafficking of lysosome-related organelles and *rab27b*, which is associated with organelle biosynthesis and maturation via the endocytic pathway^63,64^.

The hallmark of pigment cells is their ability to produce vast amounts of specific pigment molecules, which require the synthesis of purine, pteridine, or carotenoids, depending on the pigment cell type^65^. We therefore analyzed and compared the aggregated score of genes related to processes involving these pigments across the three cell populations (**Fig. 5**). As expected, iridophores exhibited higher levels of expression of genes involved in de novo purine synthesis but a lower expression of genes related to pteridine synthesis and carotenoid processing (**Fig. 5 A-C**). Conversely, xanthophores exhibited a higher level of expression of genes associated with pteridine and carotenoid synthesis but lower levels of genes for purine synthesis (**Fig. 5 A-C**).

**Figure 5.**
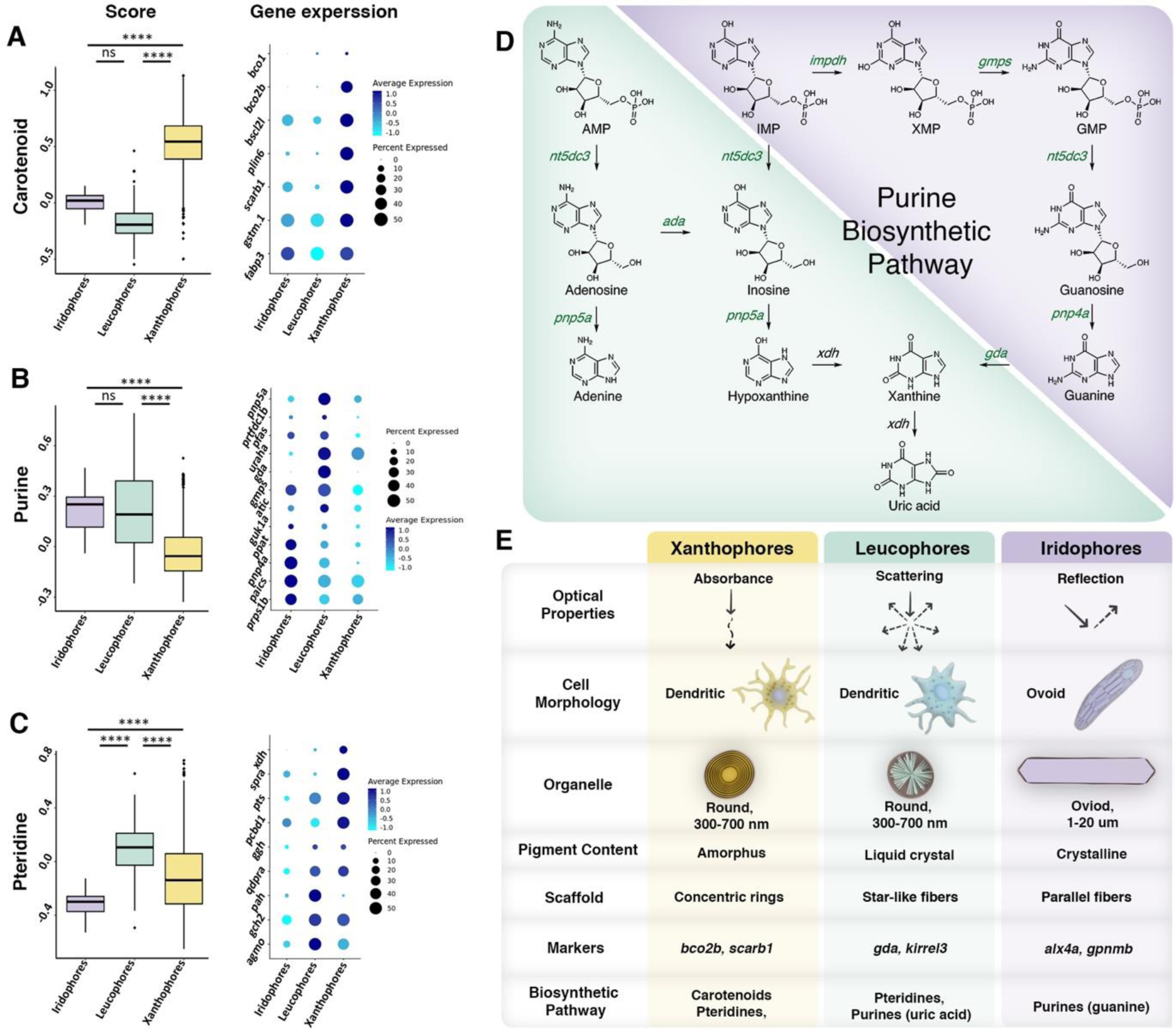
Both the purine and pteridine pathways are upregulated in leucophores. (A-C) The aggregated score for carotenoids (A), purines (B), and pteridines (C) genes calculated using the “AddModuleScore” function of Seurat (left), and the expression of the individual genes that were used in the calculation in the form of a dot plot (right). In leucophores, both purine and pteridine pathways are upregulated (Kruskal–Wallis test: purine – *X^2^*= 319.41, df= 2, p-value < 0.001; carotenoid – *X^2^*= 736.19, df= 2, p-value < 0.001; pteridine – *X^2^*= 214.02, df= 2, p-value < 0.001) followed by Dunn’s test (purine: L-X - Z=16, p<0.001; L-I - Z=1.2, p=0.59. Pteridine: L-X - Z=13, p<0.001; L-I - Z=8.8, p<0.001.) Box plots, medians with inter- quartile ranges; different letters indicate differences significant in post hoc comparison. **(D)** Schematic representation of the uric acid biosynthetic pathway, illustrating differential regulation of proteins. The selective upregulation of specific enzymes in leucophores highlights their specialized roles in uric acid biosynthesis and crystal formation. Pathways shared by both iridophores and leucophores are shown in purple, while leucophore-specific pathways are marked in turquoise. (**E**) A comparison between leucophores, iridophores, and xanthophores summarizing key observations and highlighting shared and distinct features across the three cell types.

Interestingly, leucophores displayed high levels of genes involved in both purine and pteridine synthesis, indicating either a potentially complex biosynthetic pathway or the possible production of multiple end products. To further investigate these metabolic differences, we subjected FACS sorted pigment cells to high-pressure liquid chromatography (HPLC). The results revealed that iridophores contain high levels of guanine, some hypoxanthine, and much lower levels of uric acid (**Fig. S9**), while xanthophores have high levels of various pteridines, including isoxanthopterin, L-biopterin, and sepiapterin, along with much lower levels of purines (**Fig. S10**). In contrast, leucophores contain high levels of uric acid, some adenine, and lower levels of hypoxanthine and guanine. Notably, none of the xanthophore-specific pteridines were detected in leucophores (**Fig. S11**).

To investigate the biosynthetic pathway responsible for uric acid formation in leucophores, we focused our transcriptomic analysis on mapping the purine metabolic network in both leucophores and iridophores, starting from inosine monophosphate (IMP) (**Fig. 5D**). The results revealed wide upregulation of the purine metabolic network, with some cell-specific paralog expressions, such as *pnp5a* in leucophores and *pnp4a* in iridophores (**Fig. 5D, Table S1**). However, guanine deaminase (*gda*), which converts guanine to xanthine and is downregulated in iridophores^45^, is upregulated only in leucophores (**Fig. 5D, Fig. S12**). Additionally, the relatively high expression of adenosine deaminase (*ada*) suggests that adenosine and inosine may also be converted into uric acid in leucophores (**Fig. 5D, Fig. S12**). This could also explain the relatively high levels of adenine we detect in the leucophores metabolome (**Fig. S11**).

## Discussion

A fundamental question in pigment cell biology and the study of biogenic molecular crystals is whether different crystal chemistries require distinct molecular pathways, or whether similar cellular processes can drive the formation of structurally diverse crystals. Recent investigations into the molecular mechanisms, development, and ultrastructural changes underlying guanine crystal formation^19,45-47^ have provided a foundation for comparison with the previously unexplored processes involved in uric acid crystal formation in leucophores. This comparison offers critical insights into the potential shared or divergent pathways that regulate crystal formation across different pigment cells.

Leucophores were initially believed to resemble iridophores because of their ability to form purine-based crystals^40,41^. However, recent findings suggest that the developmental and differentiation pathways of leucophores rather resemble those of xanthophores^24,42-44^. Here, we conducted detailed transcriptomic and ultrastructural studies on leucophores, xanthophores, and iridophores in order to examine how these pigment cells are related (**Fig. 5E**).

Morphologically, leucophores exhibit a similar dendritic structure to that seen in xanthophores, and both cell types contain hundreds of spherical organelles (500-1000 nm), whereas mature iridosomes in iridophores are much larger, extending up to several microns in length. However, while the pterinosomes in xanthophores contain a concentric arrangement of lipids thought to store pigment molecules, we found that leucosomes, like iridosomes, feature fibrous structures that may serve as scaffolds for purine crystal formation. Notably, unlike the parallel fiber arrangements in iridophores, the fibers in leucophores appear to radiate from the center of the leucosome to form star-shaped configurations. High-resolution microscopy and spectroscopy revealed that the uric acid within the leucosomes is arranged in a concentric assembly of nanoscopic rods projecting radially outward from the core of the sphere, and potentially aligned with the underlying fiber framework. This arrangement was recently shown to scatter light efficiently in shrimps^51^. Finding a similar structure in vertebrates suggests that these structures are more common than initially believed.

Genetically, the substantial number of upregulated genes shared by leucophores and both xanthophores and iridophores, is greater than the number of genes common to xanthophores and iridophores. Notably, genes involved in de novo purine metabolism are expressed in both leucophores and iridophores. Early enzymes in the purine biosynthetic pathway, such as *paics,* is upregulated in both cell types, suggesting a shared pathway for the conversion of phosphoribosyl pyrophosphate (PRPP) to IMP. Interestingly, downstream enzymes display much more cell-specific expression.

One key difference is the upregulation of guanine deaminase (*gda*) in leucophores but not in iridophores. This enzyme converts guanine to xanthine, a precursor for uric acid, underscoring the divergence in purine metabolism between these two cell types: guanine production in iridophores versus uric acid production in leucophores. Additionally, the selective upregulation of *nt5* and *Pnp* family members further supports the notion of cell-specific biosynthesis in each type of chromatophore. These findings support the results of our metabolome studies, which revealed differences not only in the primary crystal components but also in the minor constituents, namely hypoxanthine in the guanine crystals of iridophores and adenine in the uric acid crystals of leucophores.

Interestingly, the enzyme xanthine dehydrogenase (*xdh*), which is responsible for converting xanthine to uric acid^66^, is upregulated only in xanthophores. This could either suggest the presence of an alternative enzyme in leucophores that performs this conversion, or reflect a limitation in the sensitivity of our scRNA-seq data. Overall, these results highlight the specialized purine metabolism in each pigment cell type, reflecting their unique biosynthetic roles in pigment production.

Similarly, the findings that leucophores express high levels of genes related to pteridine biosynthesis, may suggest the presence of a complex biosynthetic pathway or the production of multiple end products. Notably, the lack of detectable pteridines in leucophores implies that these biosynthetic intermediates may be rapidly funneled into purine metabolism, and contribute to uric acid production.

The upregulation of SLC family genes in both leucophores and iridophores suggests shared mechanisms for transporting nucleobases and their derivatives. However, while some SLCs are common to both cell types, others are uniquely expressed, highlighting cell-specific adaptations for handling different molecular moieties. Additionally, genes involved in regulating the organelle micro-environment, such as V-ATPases, are upregulated in both leucophores and iridophores. Our previous work demonstrated that pH regulation within iridosomes is essential for guanine crystal formation in iridophores^67^. Given that the solubility of uric acid, like guanine, is highly pH-dependent^68^, the upregulation of similar genes in leucophores indicates that pH regulation may also play a critical role in uric acid assembly. Future studies will be necessary to determine whether pH control mechanisms in leucophores mirror those in iridophores.

Leucophores share several genes with xanthophores, including *pax7a*, a gene implicated in the differentiation of both cell types^24,61^. Additionally, *slc2a15b*, another SLC family gene upregulated in both leucophores and xanthophores, has been shown to affect the function of both cell types when impaired^43^. Our analysis now reveals that xanthophores can be subdivided into three distinct clusters based on gene expression profiles, with each displaying differential regulation of genes involved in purine, pteridine, and carotenoid biosynthesis. Cluster 1 exhibits upregulation in purine and pteridine biosynthesis pathways, while clusters 0 and 4 display upregulation of genes primarily associated with carotenoid biosynthesis (**Fig S13**). These findings suggest that different populations of xanthophores may be specialized in producing various pigments, thereby contributing to the wide range of colors, from pale yellow to deep orange, observed in xanthophores.

Perhaps the most intriguing set of upregulated genes in leucophores are those that remain unannotated. These genes may be involved in leucophore-specific processes, and some could potentially possess previously unknown biological functions. Future research focusing on selectively targeting and characterizing these genes will be crucial to advancing our understanding of their roles in leucophore cell biology.

In conclusion, our findings provide new insights into the molecular and cellular mechanisms underlying uric acid crystal formation in leucophores, revealing both shared and distinct features compared to iridophores and xanthophores. While leucophores show significant overlap with xanthophores in terms of ultrastructure and gene expression related to differentiation, the crystal formation mechanism and the biosynthetic machinery for purine metabolism resembles more closely that of iridophores, albeit with key differences. These results address the fundamental question of whether different crystal chemistries require distinct molecular pathways. We demonstrate that although leucophores share some common regulatory elements with other pigment cells, they also exhibit unique adaptations for uric acid crystal formation. This suggests that while general cellular processes can be conserved, the specific chemistry of each crystal type drives the evolution of distinct molecular pathways.

Ultimately, the results of our study highlight the evolutionary flexibility and complexity of pigment cells, and illustrate how diverse crystalline structures can arise from both shared and specialized molecular processes, thereby contributing to the remarkable diversity of coloration and optical functionality in vertebrates.

## Acknowledgments

This research was funded by an ERC Starting Grant (Grant number: 101077470, “CRYSTALCELL”) and the Israel Science Foundation (Grant No. 691/22) awarded to D.G. We extend our thanks to Prof. Kiyoshi Naruse for his support in this study and assistance in coordinating with the NBRP. We also appreciate the NBRP Medaka (https://shigen.nig.ac.jp/medaka/) for supplying multiple lines to our lab. Additionally, we would like to thank Dr. Felix Loosli for his invaluable help in establishing medaka protocols in our lab and for providing medaka fish. Moreover, we thank A. Baram for his help with the HPLC measurements and method development.

## Supplementary Information

### 1. Supplementary figure S1 - S13

**Figure S1.**
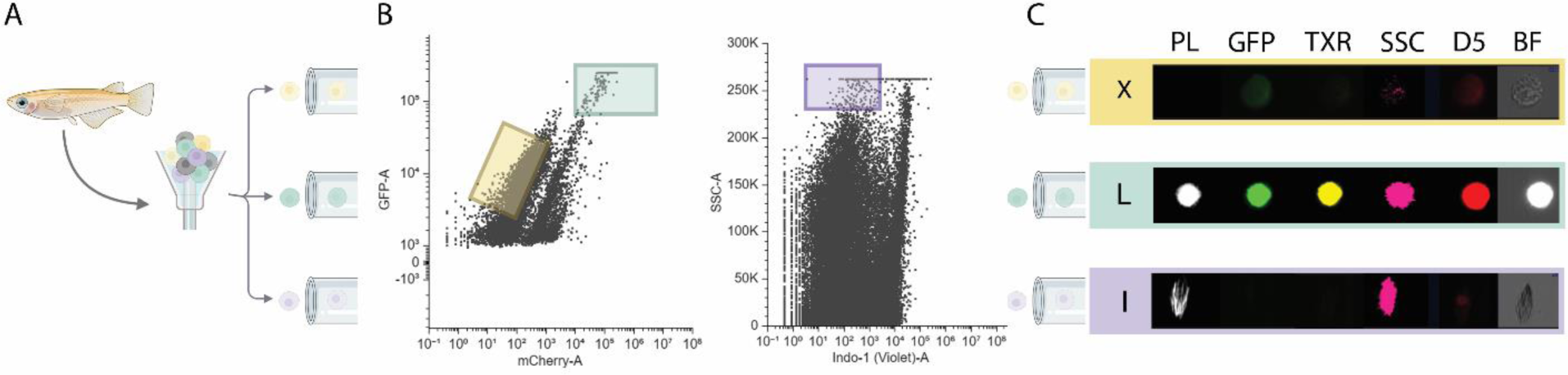
Isolation process of medaka pigment cells using fluorescence-activated cell sorting (FACS) and validation using Imagestream. The sorting was based on intrinsic autofluorescence (AF), excluding DNA-negative cells and debris. (**A**) Skin tissue and scales were carefully removed from adult HdrR medaka and dissociated into single-cell suspensions as described in the materials and methods section. (**B**) Leucophores and xanthophores were gated based on their AF in the green (GFP, 500-550 nm) and red (mCherry, 600-620 nm) channels, respectively. Iridophores were isolated using a combination of high-side scatter (SSC) and the absence of a blue signal (Indo-1, 425-475nm). (**C**) After sorting, each population was imaged using Imagestream to validate population homogeneity. Channels: Ch1 - polarized light (PL), Ch2 - GFP (500-550 nm), Ch3 - TXR (600-620 nm), Ch4 - SSC, Ch5 - Draq5 (D5), and Ch6 - bright field (BF).

**Figure S2.**
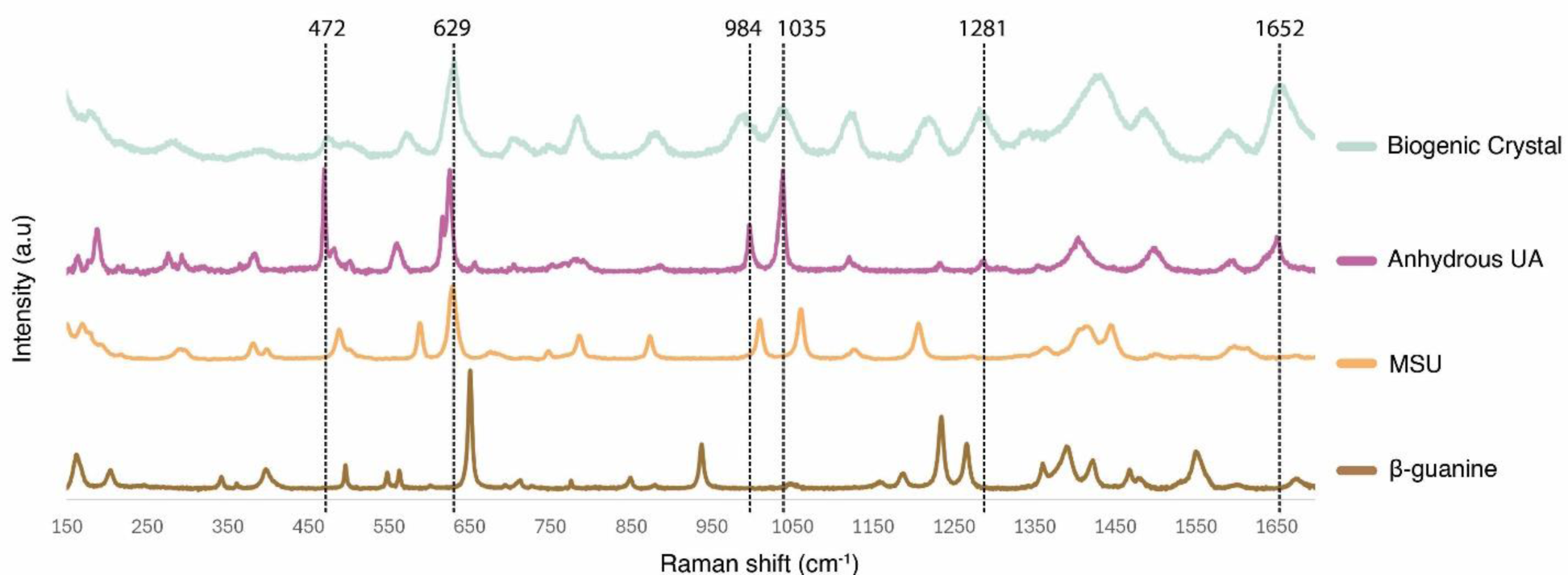
Micro-Raman spectroscopy analysis of leucophores. Raman spectra from adult medaka scales are compared with reference spectra of beta guanine (light brown), monosodium urate (MSU, yellow), and anhydrous uric acid (UA, purple). Black dashed lines mark overlapping spectral features between leucophore crystals and anhydrous uric acid powder, highlighting similarities in their molecular composition.

**Figure S3.**
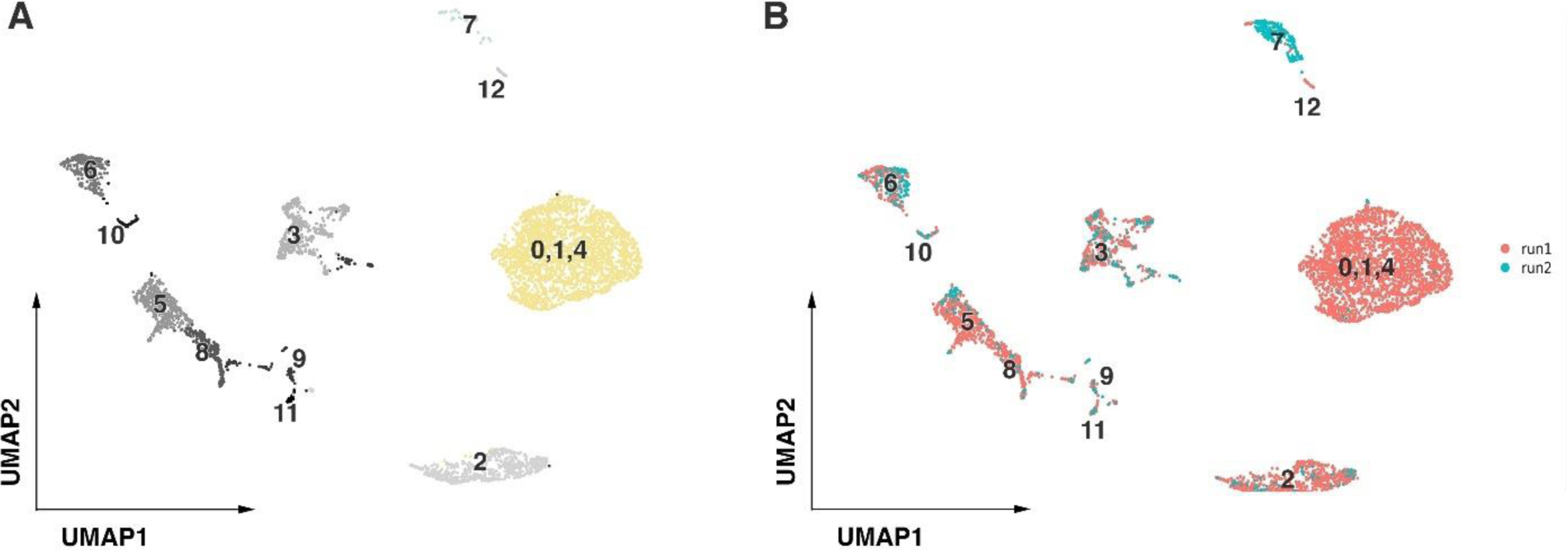
Visualization of clustered pigment cell populations in adult medaka skin tissue and scales from single-cell RNA sequencing (scRNA-seq) data following filtration. (**A**) Two-dimensional representation of scRNA-seq data from the first experiment, where clusters 0, 1, and 4 represent xanthophores (yellow), cluster 7 represents leucophores (light green), and cluster 12 represents iridophores (purple). (**B**) Comparative visualization of two separate experiments, shown as a two-dimensional reduction based on data origin. Red dots indicate cells from the first experiment as shown in panel A, while green dots represent cells from the second experiment, which focused exclusively on the collection of leucophores.

**Figure S4.**
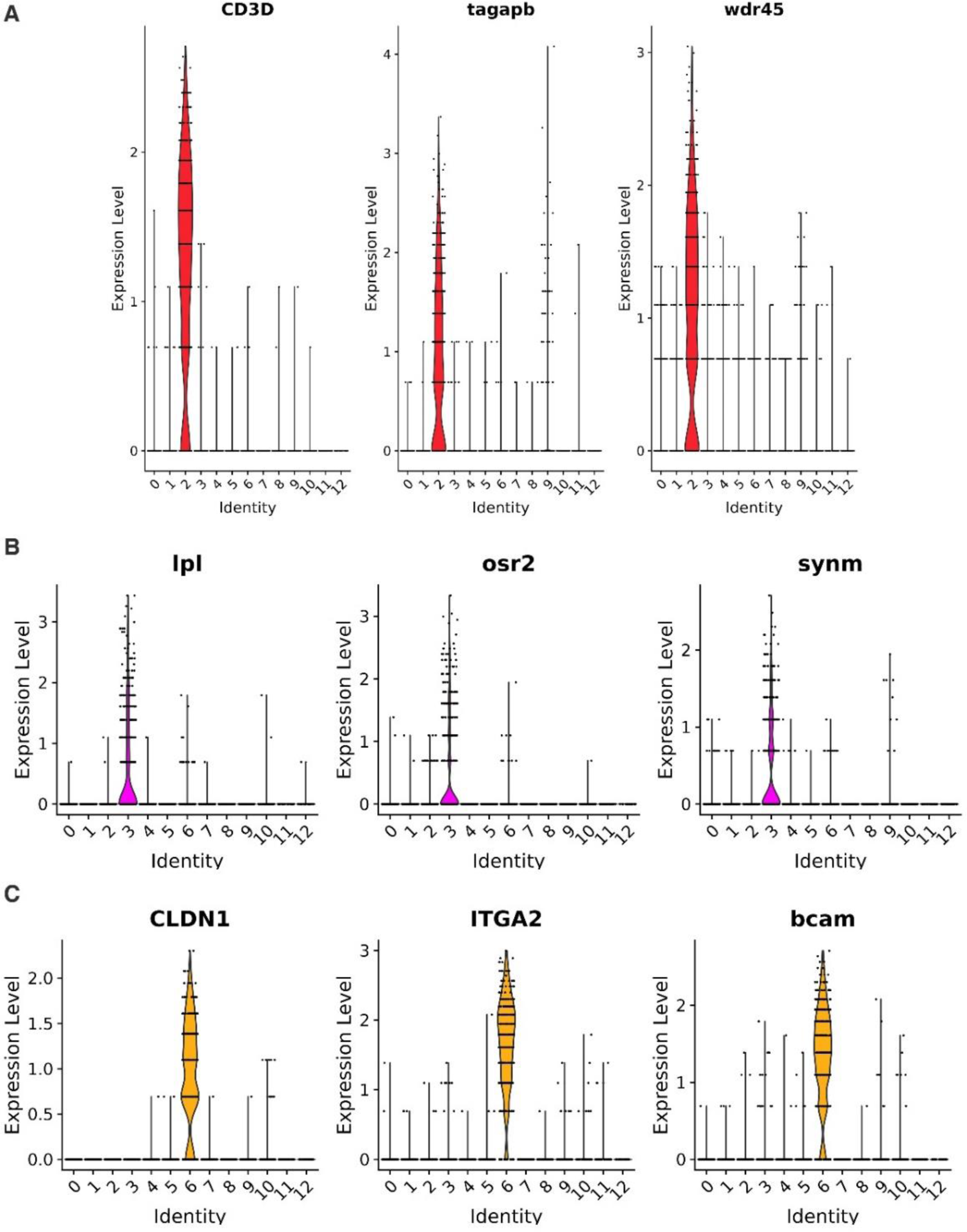
Visualization of non-pigment cell marker genes. Analysis of scRNA-seq data revealed the presence of non-pigment cells unintentionally co-isolated with pigment cell populations. (**A**) Violin plot showing the expression of immune cell marker genes^69^ in cluster 2. (**B**) Violin plot of fibroblast marker genes^69^ for cluster 3 (**C)** Violin plot of epithelial cell marker genes^69^ for cluster 6. In all plots, the Y-axis represents the average expression levels while the X-axis represents individual clusters in the dataset.

**Figure S5.**
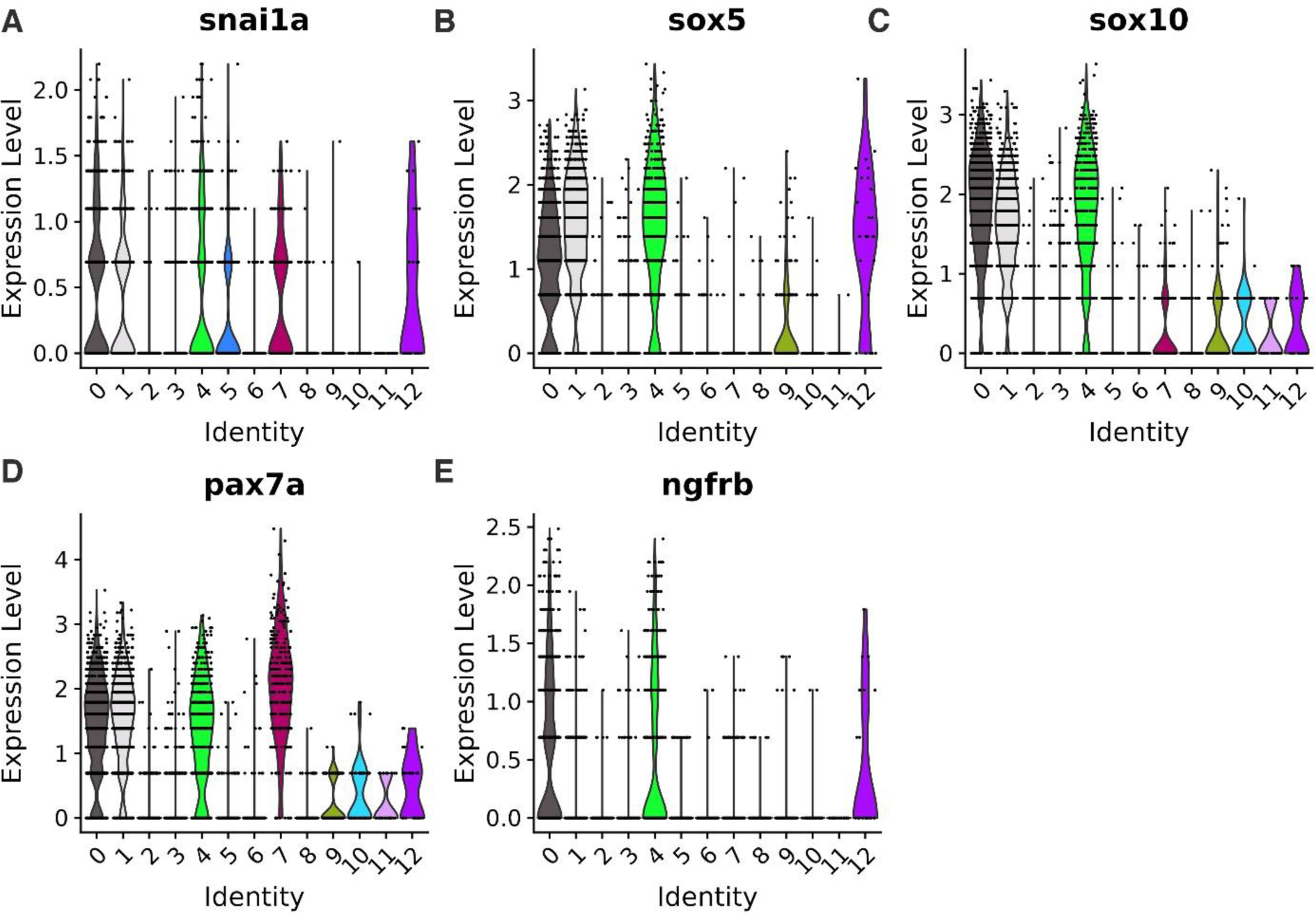
Visualization of neural crest (NC) marker genes. Detailed analysis of scRNA-seq data indicates five clusters associated with NC cells, which include the pigment cell populations. (**A-E**) Violin plots display the expression of NC marker genes, with the Y-axis representing average expression levels while the X-axis representing individual clusters in the dataset. Clusters 0, 1, 4, 7, and 12 exhibit higher expression levels of NC markers.

**Figure S6.**
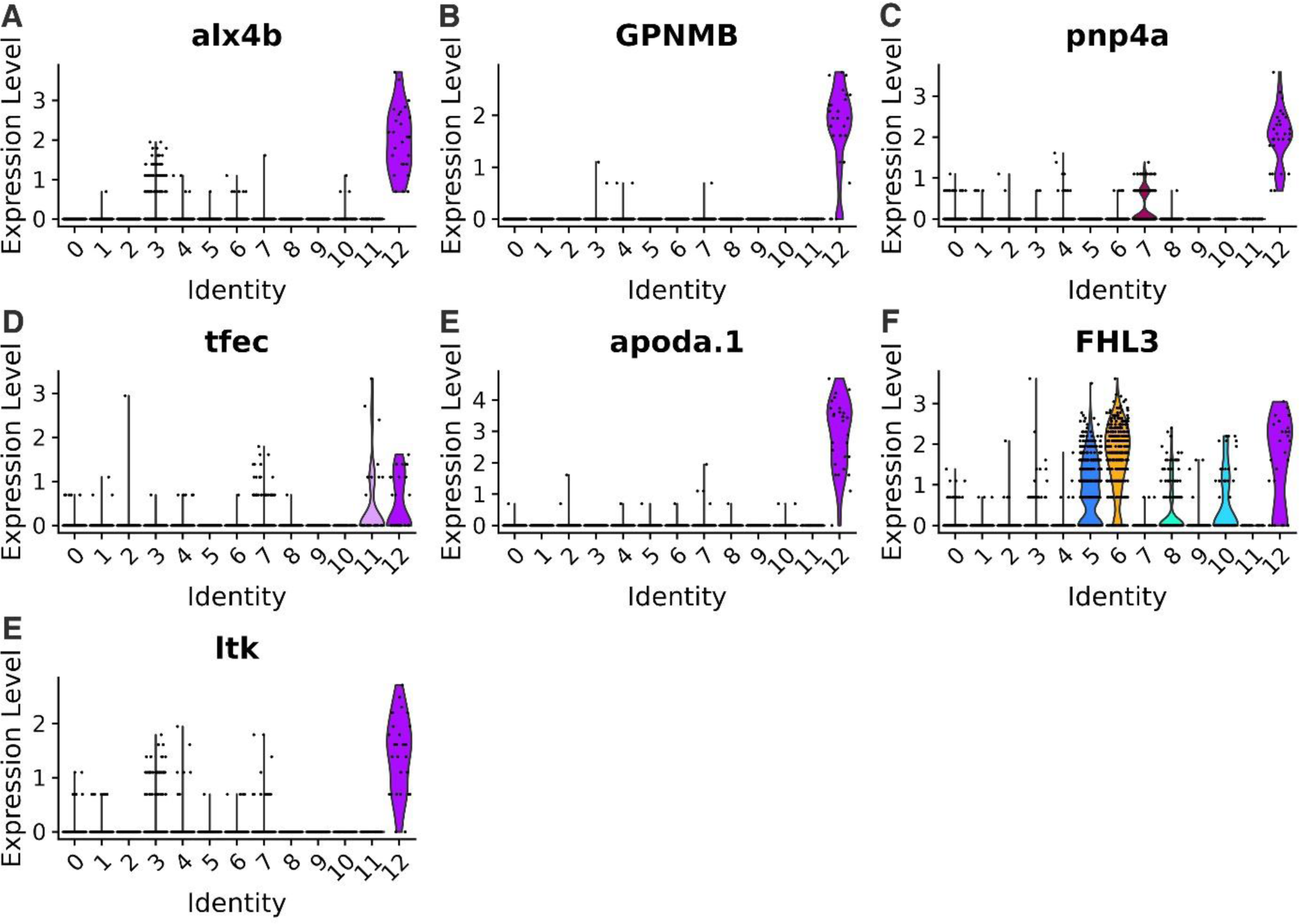
Visualization of iridophore marker genes. (**A-E**) Violin plots showing the expression of iridophore marker genes specifically within cluster 12. The Y-axis indicates the average expression levels of these markers, while the X-axis represents individual clusters in the dataset.

**Figure S7.**
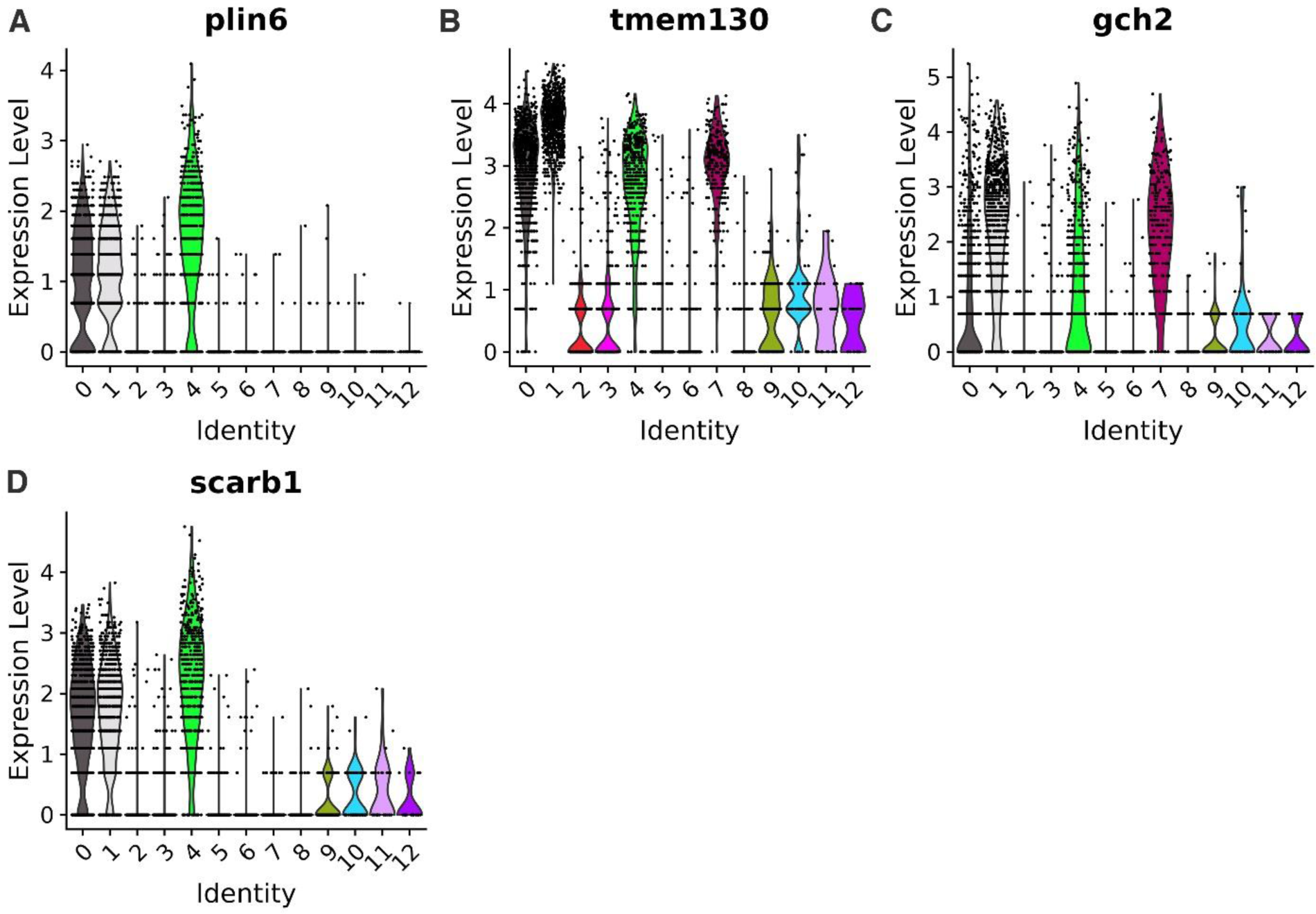
Visualization of xanthophore marker genes. (**A-D**) Violin plots displaying the expression of xanthophore marker genes within clusters 0, 1, and 4. The Y-axis indicates the average expression levels of these markers, while the X-axis represents individual clusters in the dataset.

**Figure S8.**
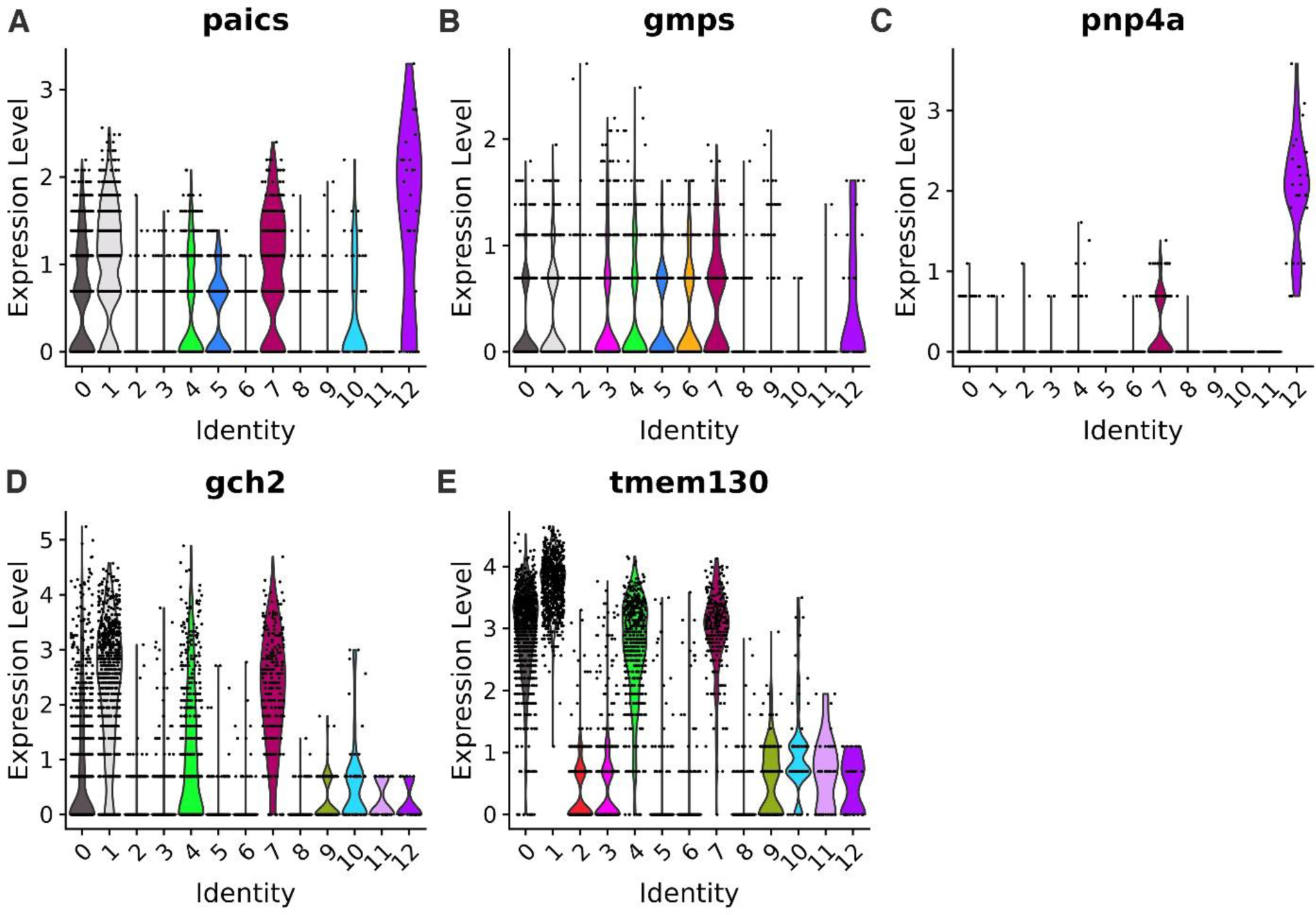
Visualization of leucophore marker genes. (**A-E**) Violin plots showing the expression of leucophore marker genes within cluster 7. The Y-axis represents the average expression levels of these markers, and the X-axis displays individual clusters across the dataset.

**Figure S9.**
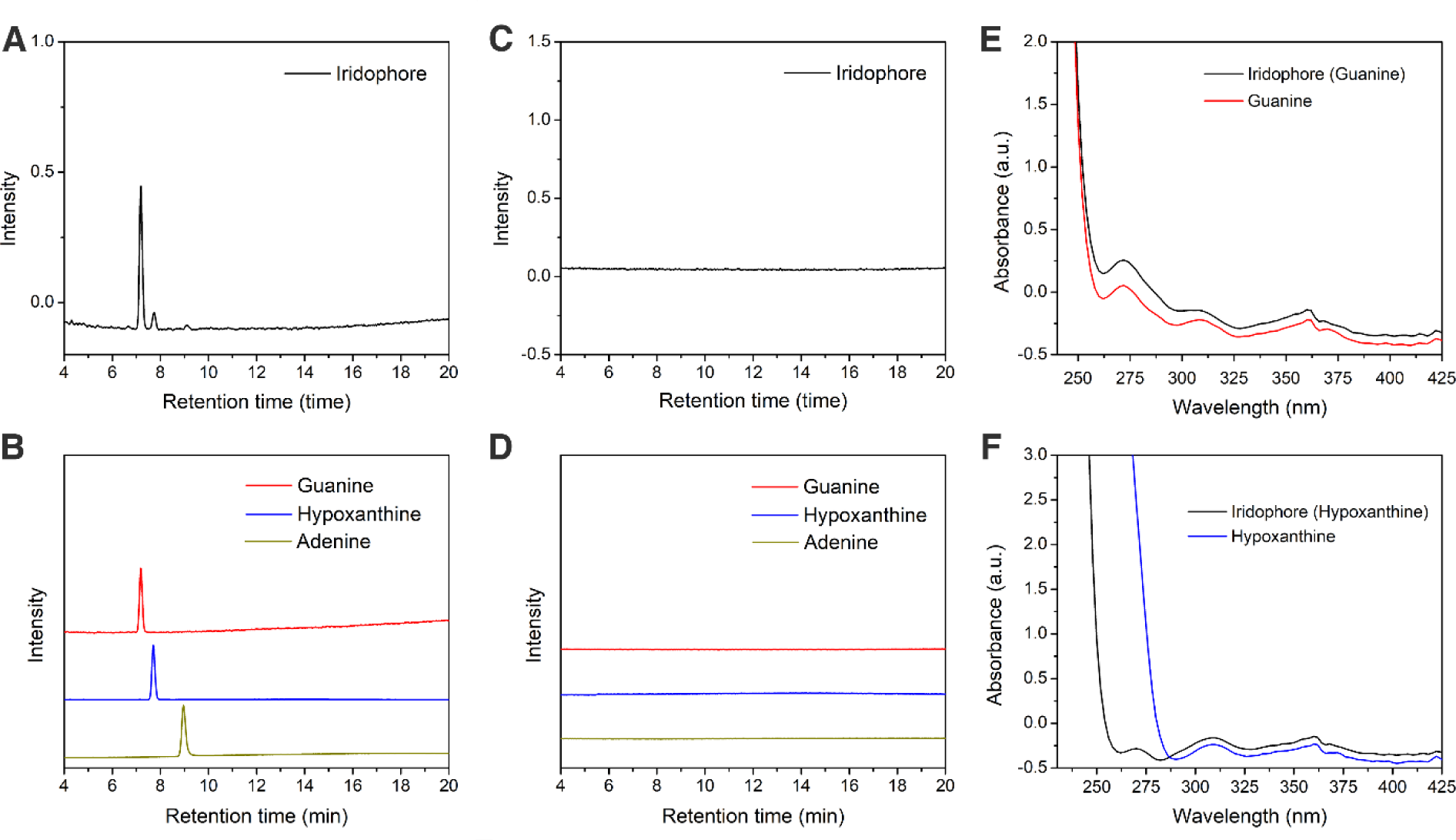
High-pressure liquid chromatography (HPLC) analysis of isolated iridophores. (**A-B**) HPLC chromatograms of isolated iridophores measured at 260 nm excitation. (**A**) Chromatogram from iridophores showing three peaks. (**B**) Chromatogram of purine standards—guanine (red line), hypoxanthine (blue line), and adenine (green line)—corresponding to the peaks observed in (A). (**C-D**) Chromatograms similar to (A-B) but measured at 340 nm excitation. (**E-F**) UV/Vis spectra representing absorbance profiles and identification of the primary compounds present in iridophores.

**Figure S10.**
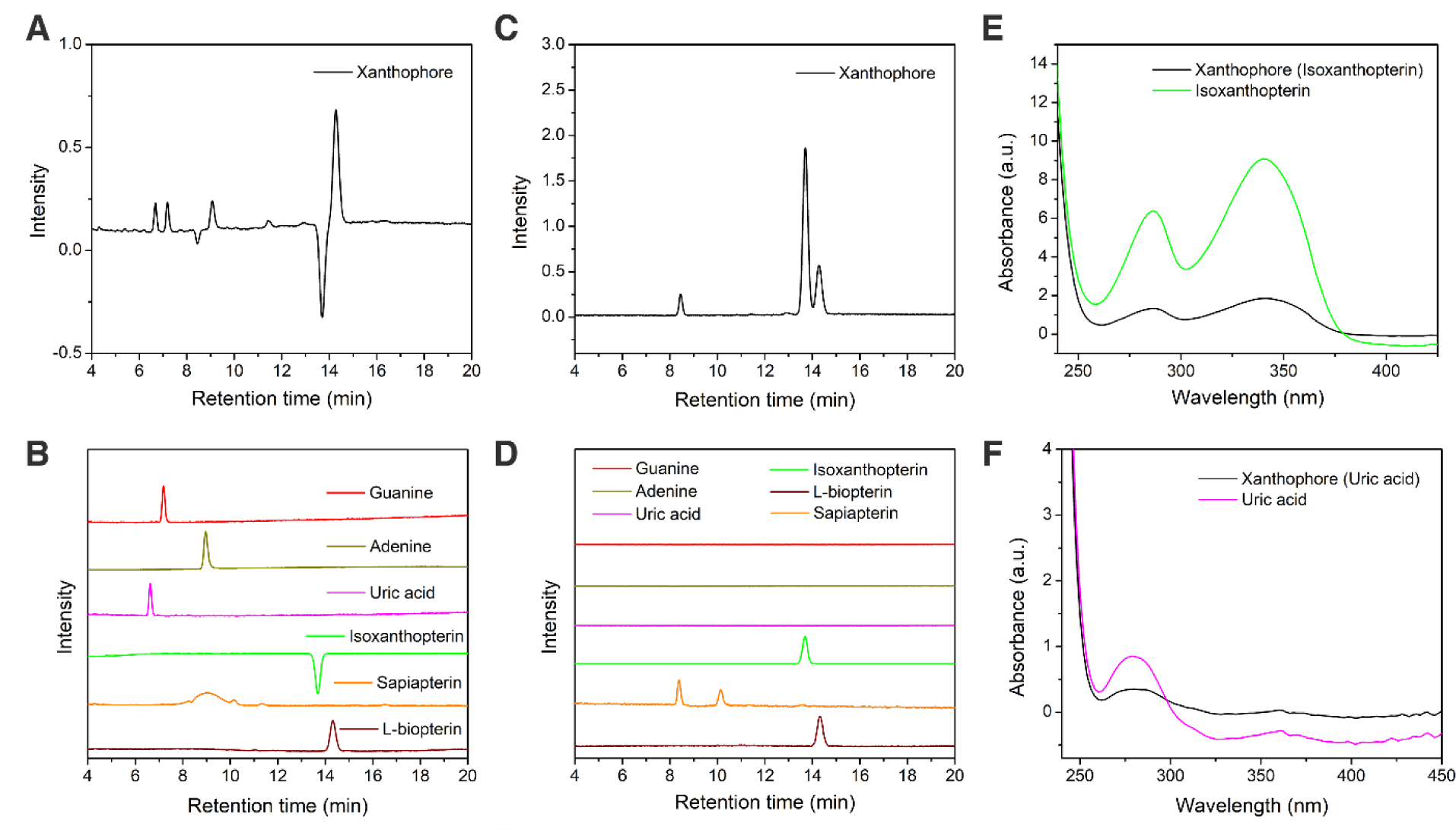
High-pressure liquid chromatography (HPLC) analysis of isolated xanthophores. (**A-B**) HPLC chromatograms of isolated xanthophores measured at 260 nm excitation. (A) Chromatogram of xanthophores showing six noticeable peaks. (B) Chromatogram of purine and pteridine standards— guanine (red line), adenine (green line), uric acid (purple line), isoxanthopterin (light green line), sepiapterin (orange line), and L-biopterin (brown line)—corresponding to the peaks observed in (A). (**C-D**) Chromatograms similar to (A-B) but measured at 340 nm excitation, which highlights the high-energy peaks of specific pteridines. (**E-F**) UV/Vis spectra indicating absorbance profiles and identification of the primary compounds present in xanthophores.

**Figure S11.**
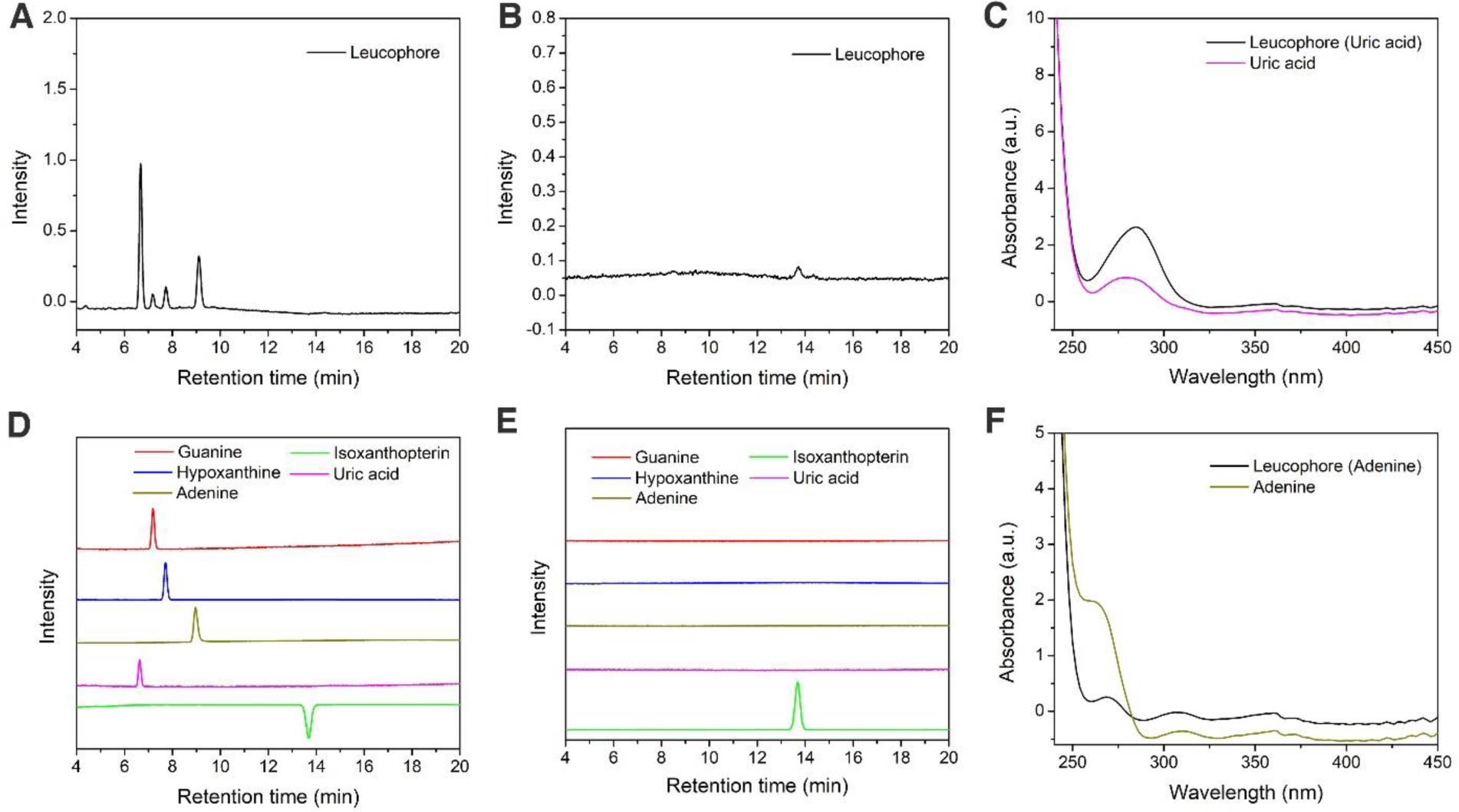
High-pressure liquid chromatography (HPLC) analysis of isolated leucophores. (**A-B**) HPLC chromatograms of isolated leucophores measured at 260 nm excitation. (A) Chromatogram of leucophores showing four noticeable peaks. (B) Chromatogram of purine and pteridine standards— guanine (red line), hypoxanthine (blue line), adenine (green line), uric acid (purple line), and isoxanthopterin (light green line)—corresponding to the peaks observed in (A). (**C-D**) Chromatograms similar to (A-B) but measured at 340 nm excitation, allowing visualization of the high-energy peak of isoxanthopterin. (**E-F**) UV/Vis spectra showing absorbance profiles and identification of the major compounds present in leucophores.

**Figure S12.**
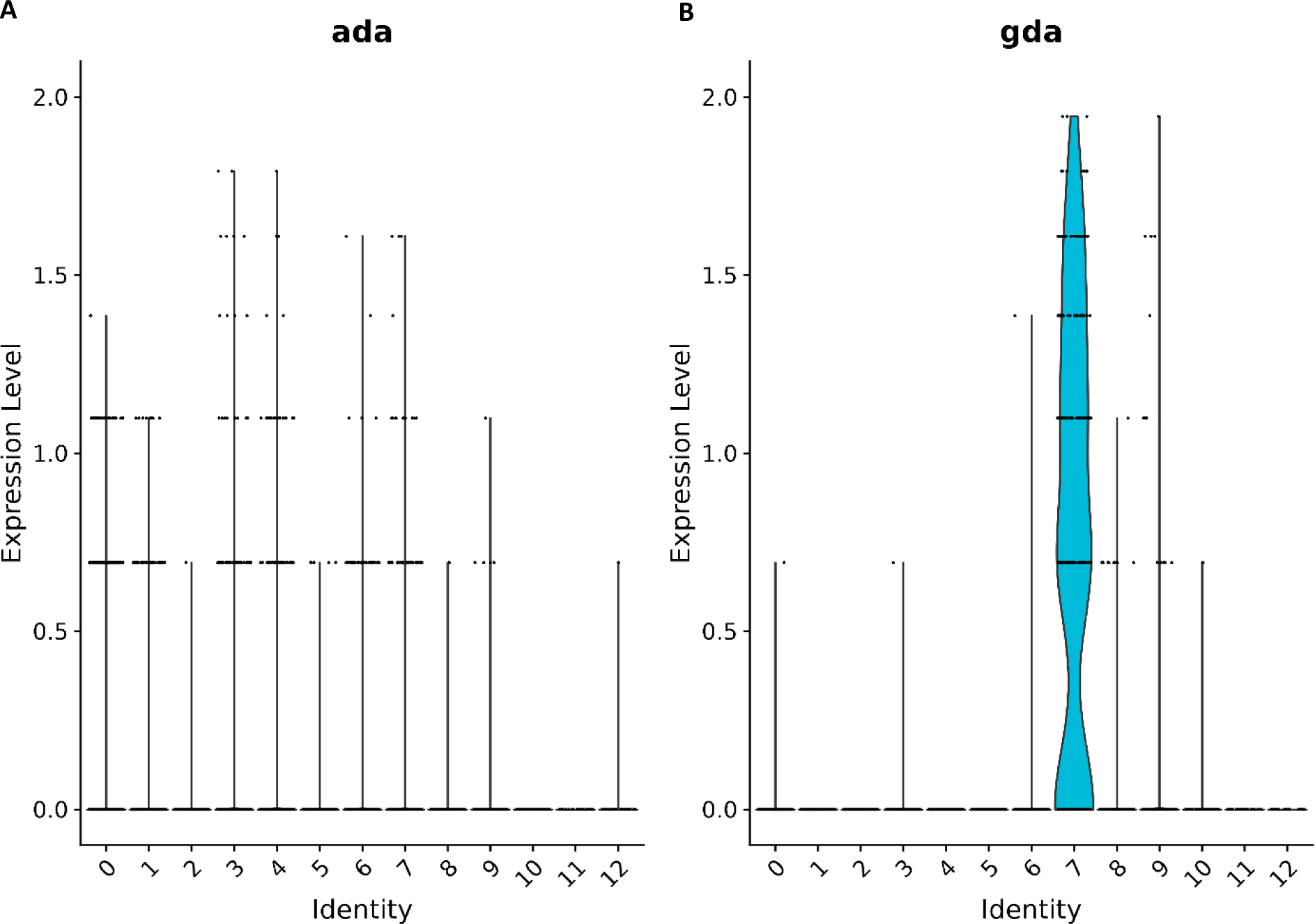
Comparative analysis of purine metabolic genes in leucophores and iridophores. (**A-B**) Violin plots highlight the expression differences in purine metabolic genes between leucophores (cluster 7) and iridophores (cluster 12). The Y-axis represents gene expression levels, while the X-axis displays individual clusters within the dataset.

**Figure S13.**
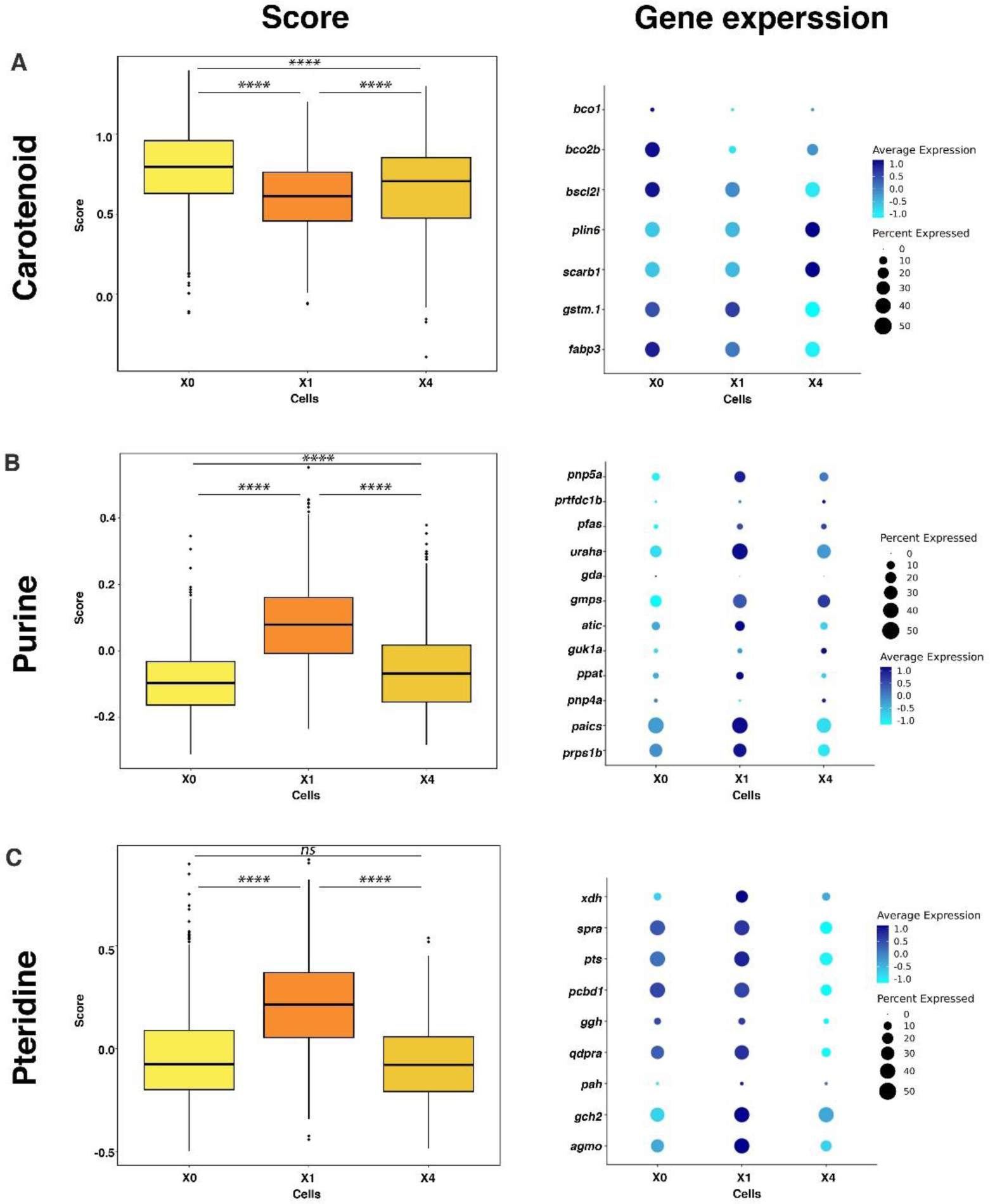
Pigmentation scoring for xanthophore subpopulations. (**A-C**) Aggregate scores for carotenoids (A), purines (B), and pteridines (C), along with corresponding dot plots showing individual gene expression within different xanthophore subpopulations (X0 = Cluster 0; X1 = Cluster 1; X4 = Cluster 4). Subpopulations X0 and X4 display similar pigmentation pathway patterns, while X1 shows a higher score for purine and pteridine pathways compared to X0 and X4. Box plots represent the score for each subpopulation, with medians and interquartile ranges. Statistical analysis was conducted using the Kruskal–Wallis test (Purine – χ² = 717.06, df = 2, p-value < 0.001; Pteridine – χ² = 525.78, df = 2, p-value < 0.001; Caretenoid - χ² = 220.87, df = 2, p-value < 0.001), followed by Dunn’s test (Purine: X1-X4, Z = 17.19, p-value < 0.001; X1-X0, Z = 26.06, p-value < 0.001,. Pteridine: X1-X4, Z = 19.27, p-value < 0.001; X1-X0, Z = 19.75, p-value < 0.001).

### 2. Materials and methods

#### Ethics and safety statement

This study followed the Weizmann Institute of Science Animal Care and Use Committee (protocol number 09331122-1).

#### Medaka husbandry and handling

Medaka (*Oryzias latipes*) were housed at ∼26°C, 14 hr light:10 hr dark and fed with Artemia and flake food. Fish were maintained and fed following standard protocols^70^.

#### Live imaging

Live imaging of cells was carried out on an inverted Zeiss LSM 900 using an Objective W Plan-Apochromat 40x/1.0 DIC M27 (FWD=2.5mm) and Plan Apochromat 63x/1.4 Oil DIC M27. Samples were illuminated with 405 nm, 488 nm, 561 nm, 647 nm laser lines. Cells were isolated via FACS or were imaged *in-situ*. After isolation of cells or hatching larvae, the sample was plated on glass-bottom well imaging plates. 0.9% low melt agarose was then added to the plates to keep cells/larva attached to the bottom for the duration of imaging.

#### Chemical fixation and Transmitted electron microscope (TEM)

Scales and larvae were fixed with 4% paraformaldehyde, 2% glutaraldehyde in 0.1 M cacodylate buffer containing 5 mM CaCl_2_ (pH 7.4) for 30 min, postfixed in 1% osmium tetroxide supplemented with 0.5% potassium hexacyanoferrate trihydrate and potassium dichromate in 0.1 M cacodylate for 1 hr, stained with 2% uranyl acetate in double distilled water for 1 hr, dehydrated in graded ethanol solutions and embedded in epoxy resin. Ultrathin sections (70–90 nm) were obtained with a Leica EMUC7 ultramicrotome and transferred to Formvar Support film slot grids (EMS). Grids were examined with a Tecnai SPIRIT transmission electron microscope (Thermo Fisher Scientific). Digital electron micrographs were acquired with a bottom-mounted Gatan OneView camera.

#### Tissue dissociation and cell isolation

Fish were anesthetized with Tricaine and euthanized on ice. Skin and scales were removed using tweezers and immersed in TrypLE Express (Invitrogen, 12604039). The sample was incubated at 37°C and shaken at 300 rpm for 1 hr, followed by mechanical disruption with a Pasteur pipette to further dissociate the cells. Cells were then strained through a 40 µm cell strainer with Dulbecco’s Phosphate Buffered Saline (DPBS) + 5% Fetal bovine serum (FBS) and centrifuged at 800 g for 10 min at 4°C. Pelleted resuspended in 5 ml DPBS + 5% FBS.

#### Fluorescence-Activated Cell Sorting (FACS)

Cells were isolated from the skin and scales of adult HdrR medaka fish as described in the ‘Tissue dissociation and cell isolation’ section and sorted using FACS. After resuspension in 5 ml DPBS with 5% FBS, the cells were stained with Draq5 (Thermo Scientific™, Cat. 62251) to label nuclei for 10 minutes before sorting. FACS analysis and sorting were conducted on a BD FACSAria II Special Order Research Product (SORP) Cell Sorter with a 100 µM nozzle, using 355 nm, 488 nm, 561 nm, and 640 nm lasers for illumination. Draq5 staining enabled the exclusion of DNA-negative debris, while additional fluorescence parameters were used to distinguish between cell populations and remove any remaining debris (see Figure S1 for more detials). Leucophores were gated based on the strong autofluorescence of their crystals, observed in the GFP (500-550 nm) and mCherry (600-620 nm) channels. Sorted cells were collected in PBS with 5% FBS and kept on ice to maintain viability.

#### Imaging flow cytometry

Cells were imaged using an Imaging flow cytometer (ImageStreamX Mark II, AMNIS corp.–part of Cytek, CA, USA). The cells were imaged using a 60X lens (NA=0.9). Lasers used were 405nm (120 mW) for Indo Blue excitation, 488nm (50mW) for AF, 642nm (50mW) for Draq5 and 785nm (1mW) for side scatter (SSC) collection. The instrument was fitted with two polarized light filters perpendicular to each other, one before the flow cell and one before the CCD camera, blocking the light completely. When a birefringent material such as crystals pass through the flow chamber, the light polarization changes and a signal is detected on Channel 1. The channels acquired were Ch01 (Polarized light), Ch02 (AF), Ch03 (AF), Ch06 (SSC), Indo-blue (Ch07), Ch09 (Brightfield), and Ch11 (Draq5). Data was analyzed using IDEAS 6.3 (Amnis Corp.). Viable cells were gated using the area and intensity of the Draq5 staining. Cells were further gated for single cells using the area and aspect ratio of the Draq5 staining, as doublets and aggregates have a larger area and lower aspect ratio. The different cell populations were identified according to the fluorescence of channels 2 and 3, and further validated using the indo-blue and polarized light channels.

#### High-Pressure Freezing of Medaka Larvae and Scales

Medaka larvae at 8 days post-fertilization (dpf) were anesthetized with Tricaine and euthanized on ice, followed by removal of the head region. Adult fish scales were carefully plucked using tweezers after anesthetization. Typically, two larvae or one scale were placed between two metal discs (2 mm in diameter; cavity size 0.2 mm for larvae and 0.025 mm for scales) in a 10% dextran solution. Samples were then cryo-immobilized using a Leica LM ICE high-pressure freezing device (Leica Microsystems, Germany).

#### Freez fracture and cryogenic scanning electron microscopy (cryo-SEM)

The high-pressure frozen Medaka larvae and scales samples were shuttled using a vacuum cryo-transfer device (VCT 100, Leica Microsystems, Germany). The sample was transferred into a freeze-etching/freeze-fracture device (BAF 60, Bal-Tec, Germany), the stage of which was maintained at -120°C and a vacuum of about 5 × 10^-7^ mbar. After fracturing, the disc remaining in the sample holder was coated with 7 nm of platinum. The coated sample was transferred to the SEM (Ultra 55, Zeiss, Germany) and observed at −120°C and a vacuum of about 5 × 10^-7^ mbar, using an acceleration voltage of 1 kV, an aperture size of 10 μm, and a working distance of 2 mm.

#### Plunge Freezing

Sorted cells (4 μL) with 15 nm gold beads (0.5 μL) were applied to glow-discharged holey carbon R2/2 Cu 200 SiO2 mesh grids (Quantifoil) coated with collagen, Type I, Rat Tail (EMD Millipore 08-115) for cell adherence. The grids were blotted and vitrified by plunging into liquid ethane using a Leica EM GP automatic plunger, under 4°C and 90% humidity conditions. Frozen grids were kept in liquid nitrogen until used.

#### CryoET

Data were collected as described in Eyal et al.^48^ using a Titan Krios TEM G3i (Thermo Fisher Scientific), equipped with a Falcon 3EC camera and a BioQuantum energy filter coupled to a K3 direct electron detector (Gatan Inc.). Data were acquired at 300 kV with the K3 camera in counting mode, using Thermo Fisher Tomography software. The TEM magnification yielded a camera pixel size of 1.6 Å (K3 counting mode), with a target defocus range of 3–8 μm. The total dose for each complete tilt series was 140 electrons per Å². Tilt series were collected with a dose-symmetric scheme from ±60° in 2° steps. Alignment and reconstruction of the tilt series images were conducted using IMOD.

#### CryoED

Samples were observed on a Talos Arctica G3 instrument (Thermo Fisher Scientific) operated at 200 kV. Images and diffraction patterns were recorded on a Falcon 3EC direct detector (Thermo Fishaer Scientific) and a OneView CMOS camera (Gatan Inc.) using imaging and diffraction modes, respectively. Analysis and processing of the movies and images were done using Digital Micrograph, GMS3 (Gatan Inc.).

#### Segmentation

Segmentation and 3D representation of the reconstructed data was done using Amira® software (Thermo Scientific). Data segmentation was performed based on contrast variations following the unique shape and structure of each component.

#### Micro-Raman Spectroscopy

Scales were placed in PBS for analysis. Uric acid and guanine reference samples were prepared using uric acid powder (Sigma Aldrich, U2625-25G) and guanine powder (Sigma Aldrich, G11950). Sample preparation followed the same method as for live imaging, with samples placed in glass-bottom well plates and immobilized using 0.9% low-melt agarose. Raman spectra were collected using a LabRAM HR Evolution (Horiba Scientific, Kyoto, Japan), a confocal Raman spectrophotometer equipped with an upright Olympus BXFM modular microscope (Tokyo, Japan). The spectrograph has an 800 mm focal length, enabling high resolution with minimal stray light, and a spatial resolution of ∼300 nm at 600 nm. The laser spot size is generally below 1 μm with a 60x objective. A LUMPlan FL N 60X WI NA=1.00 objective (Olympus, Tokyo, Japan) was used. Laser excitation was performed at 633 nm or 785 nm, with dispersive gratings of 600 or 1800 gr/mm used to optimize spectral signal-to-noise ratio (SNR). To measure fish scales, crystals were located with the microscope’s imaging capability, and spectra were obtained using both 633 nm and 785 nm lasers to reduce fluorescence. For the reference powders, individual crystal grains were selected. Spectra underwent polynomial background removal, as is standard practice.

#### Dissociation of Tissue and Preparation of Single-Cell Suspension

Tissue dissociation followed the procedure detailed in the ‘Tissue Dissociation and Cell Isolation’ section. Enriched single cells were centrifuged at 450g for 4 minutes at room temperature. The resulting cell pellet was resuspended in PBS with 0.04% Bovine Serum Albumin (BSA) and kept on ice until loading. Cell counts were determined using a hemocytometer.

#### Single-Cell Library Preparation Using the Chromium 10x Genomics Platform

Cells were counted and diluted to a final concentration in PBS with 0.04% BSA. The cell suspension was loaded onto Next GEM Chip G, targeting 10,000 cells for the first experiment and 1,000 cells for the second. The samples were processed on a Chromium X instrument to generate Gel Beads in Emulsion (GEMs, 10x Genomics). Single-cell 3’ RNA-seq libraries were prepared following the manufacturer’s protocol (10x Genomics Chromium Single Cell 3’ Reagent Kit User Guide v3.1 Chemistry).

#### Next-generation sequencing of single-cell libraries

Single-cell 3’ RNA-seq libraries were quantified using NEBNext Library Quant Kit for Illumina (NEB) and high sensitivity D1000 TapeStation (Agilent). Libraries were pooled according to targeted cell number, aiming for ∼25,000 reads per cell. Pooled libraries were sequenced on a NovaSeq 6000 instrument using an SP 100 cycles reagent kit (Illumina) with the following read length: 28 bp Read1, 10 bp I7 Index, 10 bp I5 Index and 90 bp Read2.

#### Single cell RNA sequencing bioinformatics

We obtained 4671 cells from the first experiment and 614 from the second. Alignment and generation of the count matrix were performed using the Cell Ranger (version 7.1.0) pipeline against the medaka genome (GCA_002234675.1) and using Ensemble gene annotation version ASM223467v1.110. This genome assembly, which is the most up-to-date version, is missing the sequence and the annotation of the pnp4a gene. Therefore, we added scaffold 1311 from Medaka1/oryLat2, which contains the genomic region of pnp4a, to the genome and the corresponding GTF file. Most of the downstream analyses were performed using Seurat 4.3.0. Low-quality cells, defined as cells with; >10% expression from mitochondrial genes, >1000 and < 50000 UMIs per cell and >10 or < 6000 genes, were removed from the matrix. Normalization was done using SCTransform, and cells were clustered using 20 principal components (PCs) and a resolution of 0.4. UMAP reduction was calculated using also 20 PCs. The two experiments were combined with the “merge” command of Seurat as no batch effect was observed (Fig. S3B) ^71^.

#### Venn Diagram Calculation

Differentially expressed genes (DEGs) were identified using the ‘FindAllMarkers’ function in the Seurat package in R, which applies the Wilcoxon rank-sum test to detect DEGs between cell clusters. Parameters used for this function were consistent with those in Table 1: (1) genes had to be expressed in at least 25% of cells in at least one cell population (min.pct = 0.25); and (2) a minimum log2 fold-change threshold of 0.25 (logfc.threshold = 0.25). A list of genes from three pigment cell clusters was used to create a Venn diagram, illustrating the overlap of genes among the pigment cell clusters.

#### Pigmentation Score

To calculate the pigmentation score for each cell type, pigment clusters were first separated from non-pigment clusters. Three gene sets related to purine, carotenoid, and pteridine biosynthesis, as defined by Lewis et al. ^65^, were used to calculate module scores, representing the average expression of these genes within each cell. The ‘AddModuleScore’ function was applied to calculate the average expression for each gene set (i.e., purine, carotenoid, and pteridine biosynthesis genes) per cell, and these scores were stored. Box plots were generated to visualize scores across different cell identities, and the statistical significance of differences in gene module scores between cell types was assessed using the Kruskal-Wallis test, followed by Dunn’s post-hoc test.

#### High-Pressure Liquid Chromatography (HPLC)

HPLC analysis was conducted following a customized version of a previously described ^28^. Ammonium formate buffer (10 mM, pH 3.7) was prepared by dissolving 630 mg of ammonium formate (Sigma-Aldrich, 70221-25G-F) in 1 L of double-distilled water, then adjusting the pH with ∼200 μl of formic acid (Sigma-Aldrich, 27001-1L-R), and used as mobile phase A. Prior to analysis, acid hydrolysis was performed on in vitro, biogenic, and standard samples. For in vitro samples, 1 mg of crystals was vortexed with 500 μl of perchloric acid (Sigma-Aldrich, 244252-1L) at room temperature. Biogenic crystals from isolated cell pellets post-FACS were dissolved in 120 μl of perchloric acid. Standard solutions were prepared by dissolving 90 mg of guanine, 4.5 mg of hypoxanthine, and 1 mg of adenine in 90 ml of perchloric acid. All samples were ultrasonicated for 15 minutes at room temperature. Ammonium formate buffer was used for all sample dilutions, with standard samples of guanine, hypoxanthine, and adenine diluted 5–700 times for calibration curves, while biogenic and in vitro samples were diluted at a 20:1 ratio. Before injection, samples were filtered using a 0.22 μm PVDF filter (Merck Millipore, SLGV033RS).

All HPLC measurements were performed on an Agilent 1260 Infinity system (Agilent Technologies) using an Agilent Zorbax Eclipse Plus column (250 mm × 4.5 mm, 5 μm). Optimal separation was achieved with gradient elution using phase A (10 mM ammonium formate buffer, pH 3.7) and phase B (methanol; BioLab, 1368350100) as follows: 0–10 min, 1% B; 10–13 min, linear gradient to 80% B; 13–19 min, hold at 80% B; 19–21 min, linear gradient to 1% B; 21–30 min, hold at 1% B. The total run time was 30 minutes, with a flow rate of 1 ml/min, injection volume of 10 μl, and column temperature maintained at 30 °C. Ultraviolet detection was performed at 260 nm and 340 nm.

